# Behavioral compensation preserves collective behavior when individual members are compromised

**DOI:** 10.64898/2026.03.18.712477

**Authors:** J.B. Nguyen, C.E. Lambert, C.N. Cook

## Abstract

Collective behavior in animal societies can buffer individual costs and confer resilience to environmental challenges. However, the mechanisms by which groups sustain function when members are compromised remain poorly understood. In the presented study, we investigate how social context shapes collective fanning, a thermoregulatory behavior critical for colony function, in Western honeybees (*Apis mellifera*). Using oxytetracycline (OTC), a known physiologically disruptive antibiotic to honeybees, to selectively impair certain group members, we tested our hypothesis that the presence of untreated bees would rescue the fanning response in mixed-composition groups. We show that groups containing untreated individuals fan at levels comparable to fully untreated groups, despite the presence of OTC-impaired bees. This preservation of collective thermoregulatory function was correlated with both treated and untreated individuals in mixed groups shifting their interaction dynamics and social network positions. These findings reveal a decentralized mechanism of collective resilience, whereby behavioral compensation by individuals sustains group-level thermoregulation under partial disruption. Our results provide a framework for understanding how social insect colonies maintain function in the face of individual-level perturbations, with broader implications for predicting the limits of collective resilience in animal societies experiencing increasing environmental pressures.

## Introduction

Engaging in collective behavior often allows social organisms to better deal to challenges in their environment, as it can buffer individual costs associated with responding to such challenges (Blumstein et al., 2023; Gordon, 2023; Krause et al., 2002). For example, army ants (genus *Eciton*) create sophisticated bridges with their bodies as a way for nestmates to traverse across large gaps in their environment (Garnier et al., 2013; Reid et al., 2015). Vigilant individuals such as those found in meerkat and prairie dog societies use alarm calls to warn groupmates about predators, despite increasing their own chances of predation (Hoogland, 1983, 1995; Krause et al., 2002; Townsend et al., 2012). Famously, Asian honeybee workers (*Apis cerana*) engulf invading hornets (*Vespa mandarinia*) and collectively produce heat up to 46°C to kill them (Ono et al., 1987, 1995; Sugahara et al., 2012; Yamaguchi et al., 2018). Thus, animal societies may be resilient, or able to quickly recover from disturbance, to a wide variety of environmental challenges through a decentralized sharing of costs and benefits across individuals (Reed et al., 2024; Thorogood et al., 2023).

Group resiliency is hypothesized to be a result of the variation in physiology and behavior across many individual group members (del Mar Delgado et al., 2018; Jolles et al., 2020; Júnior et al., 2022; LeBoeuf & Grozinger, 2014). In response to shifts in their surroundings, animals can adjust their behaviors and regulate their internal states to preserve homeostasis (Cannon, 1929; Davies, 2016). These behavioral and physiological adjustments vary across individuals, as seen in the variation in individual response thresholds (Abram et al., 2017; Kassahn et al., 2009; Killen et al., 2013; Koolhaas, 2008; Koolhaas et al., 2010; Pezzulo et al., 2015; Williams, 2007). Individual variation may therefore both modify and be modified by the social environment (Bentzur et al., 2021; Cook et al., 2020; Cook & Breed, 2013; Herbert-Read et al., 2013; Jäger et al., 2019; Jolles et al., 2020; Laub et al., 2024; LeBoeuf & Grozinger, 2014; Oliveira, 2009; Pinter-Wollman et al., 2011; Webster & Ward, 2011). For instance, groups comprised of at least 50% socially isolated *Drosophila* flies behave as aggressively as groups comprised of 100% socially isolated flies (Bentzur et al., 2021), and zebra finches (*Taeniopygia guttata*) preferentially sing depending on social context (Fernandez et al., 2017). Individual variation can influence communication rate in social insects (McHenry et al., 2025; Pinter-Wollman et al., 2011), and different cognitive phenotypes can modulate honeybee colony foraging by changing the enthusiasm of forager recruitment (Cook et al., 2020). Indeed, there is a nuanced relationship between individuals and the emergent collective behaviors they engage in with others in their group, and this relationship likely contributes to resiliency as an emergent property of collectives (Abram et al., 2017; Killen et al., 2013; LeBoeuf & Grozinger, 2014; Neumann & Pinter-Wollman, 2019; Pinter-Wollman et al., 2011, 2013).

However, collective resilience likely has a limit beyond which collapse can rapidly occur, and these limits are not yet well characterized. As animals experience increasing environmental challenges due to human impacts (Bradshaw & Holzapfel, 2010; Buchholz et al., 2019; Michelangeli et al., 2022; Nardone et al., 2010; Radchuk et al., 2019), there is an urgent need to identify how societies realistically balance individual homeostasis and variation while maintaining collective function. One method is to investigate group function while accounting for individual-level changes. For example, in clonal raider ant colonies (*Ooceraea biroi*), individual variation in behavior alone produces different parasitic infection rates in distinct worker castes, which can modulate overall colony social organization (Li et al., 2023). Shifts in worker demographics can trigger rapid physiological maturation to older or reversion to younger worker behavioral phenotypes to maintain colony function (Beshers et al., 2001; Huang & Robinson, 1992, 1996). Hence, characterizing the resilience of social groups under partial disruption is critical for predicting how and when collective function changes or breaks down.

As agriculturally important pollinators and eusocial insects that exhibit numerous complex behaviors, Western honeybees (*Apis mellifera*) are valuable models to understand the mechanisms of collective behavior (Davidson et al., 2021; Lemanski et al., 2019; Seeley et al., 1991; Seeley & Buhrman, 1999; Seeley & Kirk Visscher, 2004). Honeybees communicate with each other to coordinate task performance amongst themselves in several ways: the most well studied is the waggle dance, used by foragers to inform and recruit foragers to a certain resource location (Grüter & Farina, 2009; Riley et al., 2005; von Frisch, 1993). In addition to the waggle dance, honeybees perform many other tasks critical to colony survival. For instance, honeybees must regulate their colony’s internal temperature near 35°C for their brood to properly develop (Lindauer, 1955). To keep their colonies cool, middle-aged honeybees flap their wings to circulate air into the colony, a collective behavior known as fanning, and the workers that perform fanning are called fanners (Egley & Breed, 2013). Fanning depends on both temperature and social contexts, as bees experiencing high temperatures will rarely fan when isolated but will fan in groups (Cook & Breed, 2013). Fanning is also facilitated by tactile interactions, as workers that are spatially separated but can still sense each other’s presence through smell and sight are less likely to fan compared to workers that are allowed to physically touch each other (Lambert et al., 2025). The presence of an experienced fanner bee can modulate whether a non-experienced bee will fan (Kaspar et al., 2018), indicating that certain influential individuals may help maintain group function. Whether fanners can continue to coordinate fanning when some group members are compromised remains unexplored. Determining the mechanisms of the coordination of fanning in the honeybee will provide further understanding of how individual-level interactions and coordination can lead to a group-level response that is resilient to environmental challenges.

Here, we used the antibiotic oxytetracycline (OTC) as a method to impair individual honeybee fanners (Nguyen & Cook, 2025) and assess how social context shapes their behavior. OTC is used in beekeeping and is broadly known to perturb honeybee biology (Masood et al., 2022; Nguyen & Cook, 2025; Ortiz-Alvarado et al., 2020; Raymann et al., 2017), so this antibiotic is a relevant tool to study honeybee collective behavior. OTC reduces bee fanning response by decreasing the number of interactions within groups (Nguyen & Cook, 2025). In the present study, we vary the composition of groups of fanners by creating groups containing a mixture of antibiotic treated and untreated bees. We hypothesized that the presence of untreated bees would rescue the fanning response in antibiotic treated bees. We predicted that mixed groups with untreated individuals would fan similarly to uniform untreated groups, and untreated bees within these mixed groups would interact more with treated bees, influencing their fanning likelihood. By exploring these hypotheses and predictions, we provide insight into how honeybees socially organize, offer a framework to scrutinize group performance under partial disturbance, and highlight the resilience of social insects.

## Methods

All experiments described below were conducted between May and October 2022 & 2024. Honeybees (*Apis mellifera linguistica (l.)*) used in experiments originated from 9 managed colonies located on the roof of Wehr Life Sciences at Marquette University in Milwaukee, Wisconsin. Colonies were actively monitored for diseases and treated accordingly to ensure colonies remained healthy. These managed colonies have never been treated with oxytetracycline, so the bees we used in this study were naïve to the antibiotic treatment.

### Fanner collection

To collect fanners, we followed the protocols outlined in our previous work (Cook & Breed, 2013; Nguyen & Cook, 2025). Briefly, we collected actively fanning workers at the entrance of our managed colonies using forceps and grabbing their legs (Cook & Breed, 2013; Nguyen & Cook, 2025). Although age plays a role in regulating honeybee worker task (Egley & Breed, 2013; Huang & Robinson, 1996), we did not explicitly control for age in all our experiments, as we selected bees based on behavior alone. Fanners were collected into empty pipette tip boxes with 96 well plates taped inside in groups of 25-30 individuals, and each box contained fanners from the same colony. On collection days, three boxes of fanners (one box per treatment group as later described) were collected from a single colony. When enough fanners were available, an additional triplicate set of boxes were collected from an additional colony on the same day to maximize number of experimental replicates performed per week.

### Experimental treatment

To disrupt individual function, after collection, all boxes of fanners were randomly assigned one of 3 experimental treatment groups: sugar syrup solution only (referred to as “Lab-Control”) sugar syrup solution for 4 days followed by 24 hours of sugar syrup solution with antibiotic (referred to as “1-Day”), or sugar syrup solution with antibiotic for 5 days (referred to as “5-Day”). Sugar syrup solution was made to be 40% concentration, while the antibiotic treatment was 40% concentration sugar syrup solution with 0.37 mg/mL of oxytetracycline hydrochloride (Oxytet Soluble), which was procured through a local veterinarian. This treatment regimen alters honeybee health (Raymann et al., 2017, 2018) and impacts honeybee social interactions (Nguyen & Cook, 2025). To mimic internal colony conditions, bees were given treatment *ad libitum* and kept in an incubator set at 35°C for 5 days. Dead bees were removed daily from boxes.

### Fanning assays

To test fanning under varying social contexts after the treatment period, we subjected all surviving fanners to a fanning assay, following protocols outlined in our previous work (Cook & Breed, 2013; Nguyen & Cook, 2025) with a few adjustments. First, to identify individual bees, we marked bees on their thorax with a unique color based on their treatment group using non-toxic, water-based acrylic paint markers (Montana). Bees were then placed in small cylindrical wire mesh cages (340.47 cm^3^) in groups of 10 in our fanning assay set up. Bees were placed in either “uniform” groups (10 bees from the same treatment) or “mixed” groups (5 bees from one treatment and 5 bees from another treatment), creating a factorial experimental design (Figure 1). We marked all bees regardless of group composition to control for any handling stress caused by marking their thorax. Bee groups always consisted of individuals from the same colony.

**Figure 1.**
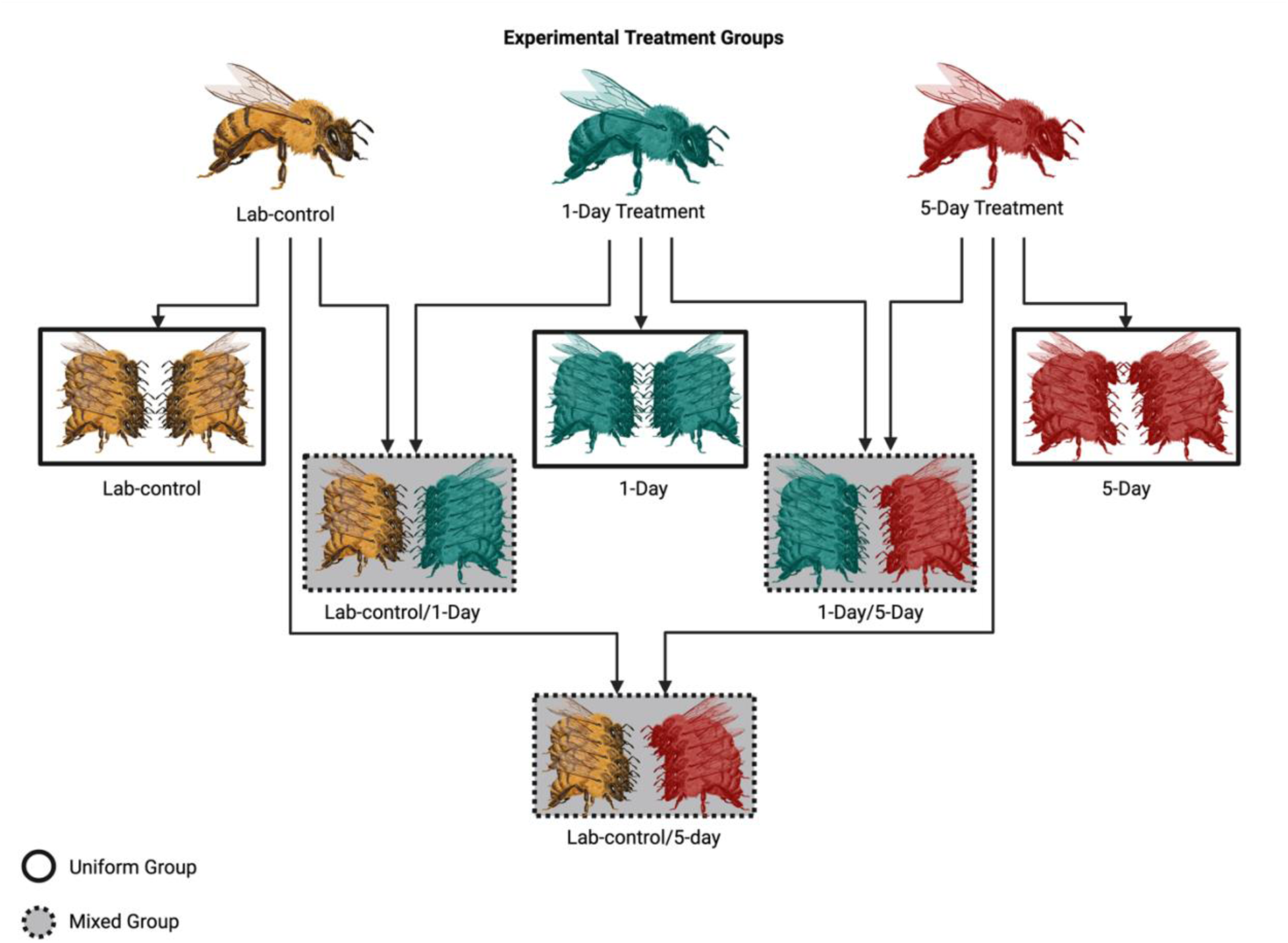
Composition of experimental mixed and uniform groups in a factorial design. Bees were placed in groups of 10 of their own treatment group (uniform) or a mixed group of 5 bees from their own treatment and 5 from another treatment (mixed). Individuals were always paired with other bees collected from the same colony. Following this factorial design, we had a total of 6 different experimental treatment groups. Solid black outlines indicate uniform groups, while dotted outlines indicate mixed groups. This color and outlining scheme are repeated for all subsequent figures. Figure was created in Biorender. Honeybee illustrations were created by Impact Media Lab.

During the fanning assay set up, we observed multiple instances in which individuals would fight (defined as actively biting and stinging another bee and releasing alarm pheromone) when placed into cages together, despite originating from the same colony. We tested whether these fighting instances were caused by our treatment and/or group composition. Because we found no statistical significance in fighting instances across group composition (Figure 2; X^2^ = 1.6589, p = 0.894), we determined that the fighting was not a result of disruption of nestmate recognition and could have been an artifact of handling the bees. Regardless, we removed and replaced bees who fought during the experimental set up to reduce confounding effects.

**Figure 2.**
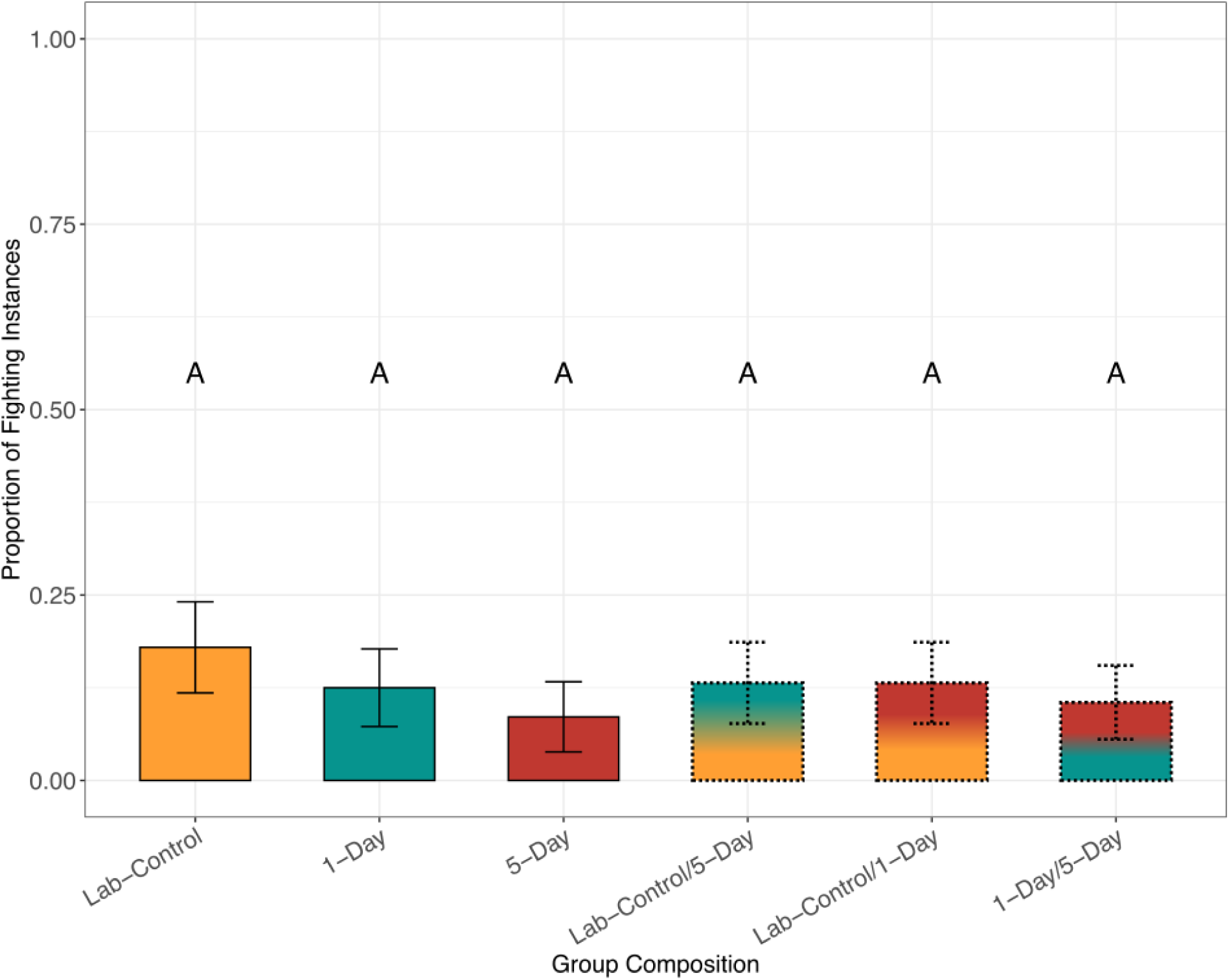
Observed fighting instances were not statistically different across group composition. During the creation of fanning groups as described in previous figure, we observed several fighting instances between individuals. We therefore recorded fighting instances as when a pair or more bees immediately grabbed onto each other and engaged in stinging, biting, alarm pheromone release, etc., after an individual was added to a cage. Out of 228 cages, only 29 cages fought, indicating that any potential novelty, treatment effect, or group composition alone did not explain the fighting behaviors we saw, and we reasoned that fighting was due to stress from handling. Any individuals that engaged in fighting were removed from trials regardless to avoid any confounding behavioral data. GLM with a binomial distribution, X^2^ = 1.6589, p = 0.894. Error bars indicate standard error.

All groups were allowed to acclimate at room temperature (mean temperature 23.42 ± 1.73°C) in 1 gallon glass jars (Uline) on hotplates (OHause Guardian 5000, model number e-G51Hp07C) with a temperature probe (Campbell Scientific, 109 Temperature Probe) wired through a GRANITE Volt 116 16-/32-Channel 5V analog input module connected to a computer. We used SURVEYOR software (Campbell Scientific, v.101) to monitor and collect temperature data in real time. To evaluate fanning, we heated the air temperature in the jar via the hotplate at 1°C/min. Any trials with ramp rates that were not within 0.1°C of this ramp rate were removed from further analysis, as heating rate has been previously demonstrated to impact fanning likelihood (Cook et al., 2016). We recorded the time, temperature, and number of fanners seen fanning throughout the assay. Fanning frequently occurs in ‘bouts’; therefore, we recorded the first group of bees fanning within 30 seconds of the first fanner as part of the ‘first fanning bout’ and the maximum number of bees seen fanning within 30 seconds of each other in the trial as part of the ‘max fanning bout’, as done in our previous studies (Cook & Breed, 2013; Nguyen & Cook, 2025). Fanners were observed until mortality, of which time and temperature were also recorded.

### Video Recording of Group Dynamics

As prior research has shown that bees treated with antibiotics moved more and have longer interactions compared to untreated, we aimed to determine how these behavioral changes impact communication dynamics of the group (Nguyen & Cook, 2025). To test this, we recorded fanning trials, used tracking software to detect interactions, and finally configured social networks.

We first video recorded bees during fanning assays. Bees were marked and placed in uniform or mixed groups in a factorial manner as described earlier (Figure 1). This box was then placed into a Pyrex baking dish (33 x 21.6 x 8.25 cm) with a glass lid, and bees were allowed to acclimate for 25 minutes. Bees were recorded using a Panasonic HC-V800 camera, and experimenters observed bees using a computer monitor connected to the camera. We began video recording, which had a 30 frames per second resolution, 5 minutes before the trial started. We then carried out fanning assays as previously described until mortality, and video recording was immediately halted after the trial was completed. As before, any trials that were not within 0.1°C of our 1°C ramp rate (Cook et al., 2016). All videos were backed up on an external hard drive (LaCie Rugged Mini 5TB) and transferred to a computer for further processing.

### Interaction Detection

To detect interactions between individuals, we first aligned the trials to the first instance of fanning, then created 5-minute trims of all videos: 4 minutes before the first instance of fanning and continuing 1 minute after. We did this to account for variation in the time and initial fanning temperature in videos. We then backed up all video trims onto an external hard drive and Marquette University’s cloud-based Microsoft OneDrive service for long-term storage.

To process our trims, we used the automated tracking software ABC Tracker to track individual level movements at 30 fps (Rice et al., 2020). Within the application, we then screened and manually corrected errors in the detection and tracking process (Rice et al., 2020). The program returns centroid coordinates for the body and head of every individual in each frame. Based on the center coordinates, body shape is manually set in the graphical application as a rectangle tightly fit around the bee body, along with a smaller polygon for head shape. Using these estimates of bee shape and tracked body center location, we used R to detect interactions. Interactions were defined as periods lasting at least 1s and no longer than 60s where the individuals’ coordinate areas overlapped (Lambert et al., 2025; Nguyen & Cook, 2025; Wild et al., 2021).

### Social Network Configuration

We configured social networks based on the tactile interactions between individuals, and compared network centrality measures using the R package ‘igraph’ (Csárdi et al., 2025; Csardi & Nepusz, 2006). Each node represents an individual bee, and edges are interactions between bees, weighted by total number of interactions (Kay et al., 2023; Pinter-Wollman et al., 2011; Richardson et al., 2022; Swain et al., 2022). We then calculated strength, closeness, and betweenness centrality metrics to interpret and assess differences in social positionality in all our networks (See Table 1; (Turetsky et al., 2020)). To obtain a more interpretable metric of closeness, we specifically calculated harmonic closeness, which is the inverse of the average shortest distance between a given bee and all other bee in the network (Farine & Whitehead, 2015; Zhang & Luo, 2017).

**Table 1.** Common social network centrality metrics and their definitions interpreted in our study.

| Centrality Metric | Definition | Biological Interpretation | Citation |
| --- | --- | --- | --- |
| Strength | Sum of weights of all edges for a given node in a network | Number of all interactions for a focal bee in a network | (Farine & Whitehead, 2015; Makagon et al., 2012) |
| Closeness | The average shortest distance from each vertex to each other vertex | Metric that measures how quickly an individual bee can reach all other bees in a network. Metric highlights bees that have a more central position within the network and therefore have more access to others. | (Farine & Whitehead, 2015; Zhang & Luo, 2017) |
| Betweenness | The number of shortest paths that flow through a node. | Metric that highlights bees that have a greater tendency to change groups than others, potentially acting as a facilitator of communication. | (Farine & Whitehead, 2015; Zhang & Luo, 2017) |

### Statistical analysis: General

All statistical analyses were performed in R (version 4.3.2) and RStudio (version 2024.12.0+467). We used the following R packages for our analyses: “dplyr”, “emmeans”, “fitdistrplus”, “FSA”, “ggplot2”, “igraph,” “lme4”, “tidyr”, “patchwork”, and “performance”. For all generalized linear models (GLMs), we verified model parameters and fit using the “check_model” function in the “performance” package before continuing with analyses.

### Statistical analysis: Assessing changes in fanning behavior

To test for differences in fanning behavior, we created several generalized linear mixed models (GLMMs). We first calculated the proportion of fanners in the first fanning bout and max fanning bout by dividing the number of fanners seen in each bout by the total number of bees in the group. We then created GLMMs with a binomial distribution (family = binomial(link = “logit”)) with proportion of fanners as our response variable, group composition as our predictor variable, and colony as a random effect since we sampled from 9 different colonies. We also tested whether fanning likelihood was different depending on group composition type (mixed or uniform) by creating a binomial GLMM with an interaction effect between individual treatment and group type with proportion of fanners as our response variable. We tested all models using an ANOVA Type II Wald Chi-Square test. We then extracted estimated marginal means using the “emmeans” package in R to compare individual predicted fanning probabilities depending on group composition type. Colony origin accounted for (variance ± standard deviation) 0.1514 ± 0.3891 of the first fan bout proportion, 0.6207 ± 0.7879 of in max fan bout proportion, and 0.5192 ± 0.7206 of the group composition type.

To test for statistical differences of the temperature thresholds these fanning bouts occurred, we created linear mixed models (LMMs) with temperature as our response variable, group composition as our predictor variable, and colony as a random effect. Colony origin accounted for 17.34 ± 4.164 of the variation seen in the first fan bout temperature degree and 8.863 ± 2.977 of the variation seen in the max fan bout temperature degree. All temperature threshold models were tested using an ANOVA Type II Wald Chi-Square test.

### Statistical analysis: Assessing changes in group dynamics

To test for differences in average group movement velocity and head-to-head interactions across group composition, we followed the analysis methods to quantify interactions and calculate group velocity as outlined in our previous study (Nguyen & Cook, 2025). After quantifying head-to-head interactions per bee, we created a GLM with the total number of head-to-head interactions as our response variable and group composition as our predictor variable. Then, to assess any differences in interaction dynamics in mixed groups, we calculated the expected proportion of mixed interaction pairs and uniform interaction pairs based on combinatorics of a group of 10 bees made up of 2 treatment groups followed by a Chi-square test comparing these expectations to our observed proportions. For average group velocity, we created a GLM with a gamma distribution (family=Gamma(link = “inverse”)) with average group velocity as our response variable and group composition and social context as our predictor variables.

We then tested all models after verifying model parameters using an ANOVA Type II Wald Chi-Square test and a Tukey’s post-hoc test when appropriate. We also compared the individual velocities of bees within mixed group contexts by filtering the data by treatment identity and creating new GLMs with a Gamma distribution with individual velocity as our response variable and our predictor variable to be partner treatment identity (i.e., the other treatment group mixed with the focal individual in the mixed groups). To compare average interaction durations across groups, we used a Kruskal-Wallis Test followed by a post-hoc Dunn’s Test. To test for differences in individual social position in social networks, we compared all individual centrality metrics (i.e., strength, closeness, and betweenness) across all treatment groups with different group compositions via Kruskal-Wallis Tests followed by a post-hoc Dunn’s Test with a Holm’s adjustment when appropriate.

To assess differences in positionality within social networks, we created linear models with a log-normal distribution and each of our centrality metrics as the response variable and group context as a predictor variable. We tested all models using an ANOVA Type II Wald’s Chi-Square test followed by a post-hoc comparison when appropriate. We adjusted all p-values using a Holm’s adjustment as we did not have a uniform number of videos recorded and thus social networks calculated per group.

## Results

### Presence of Lab-Control bees rescues overall fanning performance in groups of 10

We found that group composition had a significant effect on the proportion of fanners in the max fanning bout (Figure 3a, X^2^ = 43.009, p < 0.0001). Specifically, uniform 5-Day groups fanned significantly less than uniform Lab-Control, uniform 1-Day, mixed Lab-Control/1-Day, and mixed Lab-Control/5-Day groups during the max fanning bout (Figure 3a; Supplementary Table 1.2; Tukey’s post hoc, p < 0.001). In other words, fanner groups where all individuals are perturbed by antibiotic treatment for 5-days had significantly lower fanning performance compared to all other groups. Interestingly, these uniform 5-Day groups fanned similarly to mixed 1-Day/5-Day groups (Figure 3a; Tukey’s post hoc, p =0.1553), indicating that even with group members with partial disruption (1-Day treatment), group context was not enough to rescue fanning in these groups. Mixed Lab-Control/5-Day and mixed Lab-Control/1-Day groups fanned similarly to uniform Lab-Control and uniform 1-Day groups (Figure 3a, Tukey’s post hoc, p > 0.05), and mixed 1-Day/5-Day groups fanned significantly less than uniform Lab-Control and mixed Lab-Control/5-Day groups (Figure 3a, Tukey’s post hoc, p = 0.0359). We found no effect of group composition on the proportion of fanners in the first fanning bout (Supplemental Figure 2, X^2^ = 2.6567, p = 0.7527) or the first fanning bout temperature threshold (Supplemental Figure 2, X^2^ = 2.5045, p = 0.7758). There was also no effect of group composition on the max fan bout temperature threshold (Figure 3b, X^2^ = 6.2517, p = 0.2825). Overall, these results show that social context significantly impacted overall group fanning performance.

**Figure 3.**
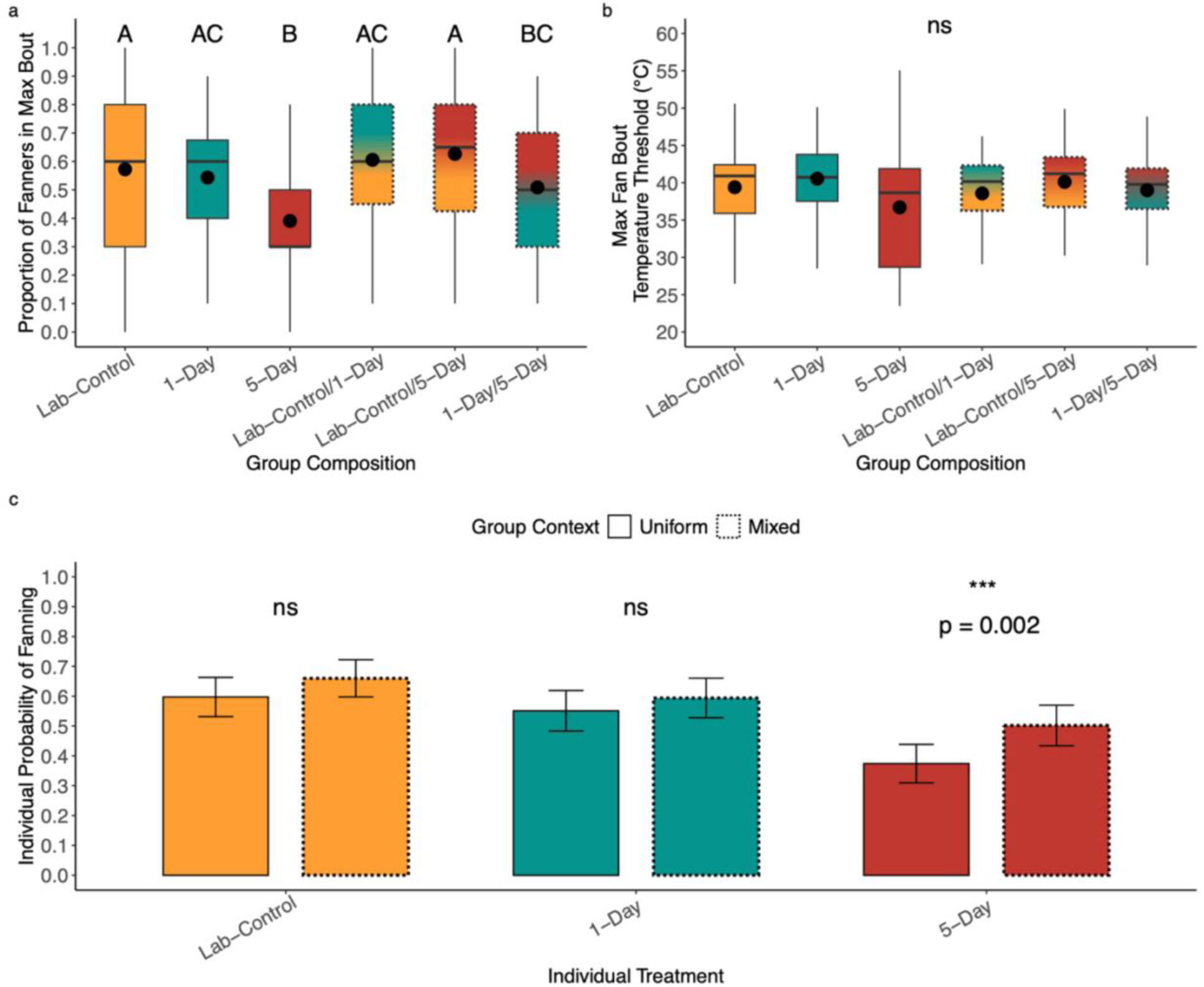
Presence of Lab-Control bees rescues collective function and individual fanning likelihood in mixed groups. N_Lab-Control_ = 38, n_1-Day_ = 39, n_5-Day_ = 35, n_Lab-Control/1-Day_ = 36, n_Lab-Control/5-Day_ = 40, n_1-Day/5-Day_ = 38, for n = cages of 10 bees. Max bout refers to the maximum number of fanners seen fanning during a trial a) Proportion of fanners in the max fanning bout seen in trials. Uniform 5-Day bee groups significantly fanned less than Lab-Control, 1-Day, 5-Day, Lab-Control/1-Day, and Lab-Control/5-Day groups but fanned similarly to 1-Day/5-Day groups. GLM with a binomial distribution, X^2^ = 43.009, p < 0.0001; Tukey’s post-hoc tests, p < 0.001 b) Temperature at which the max fanning bout of bees was seen fanning. No effect of group composition on the thermal threshold. LM, X^2^= 6.2517, p = 0.2825. Dots in boxplots represent mean while horizontal lines represent median. Letters indicate statistical significance, where groups that share letters are not statistically different from each other while groups with different letters are statistically different as determined via Tukey post-hoc test c) Probability of an individual to fan based on group context and the treatment the individual received. 5-Day bees were more likely to fan when they were in mixed groups compared to when they were in uniform groups. Estimated marginal means contrast, z-ratio = 3.060, p-value = 0.0022. Graph is plotted with the estimated marginal means probability calculated from our max fan bout logistic regression model. See Supplementary Table 1.3 for specific outputs. Error bars represent standard error. ns = no significant difference.

We found that in groups of 10 the individual fanning probability for 5-Day bees was significantly higher in mixed groups compared to uniform groups in groups of 10 (Figure 3c, Supplementary Table 1.3; *z*-ratio = 3.060, p-value = 0.0022). Individual 5-Day bees were 50.2% likely to fan when in mixed groups and only 37.4% likely to fan in uniform groups (Figure 3c; *z*-ratio = 3.060, p-value = 0.0022). In contrast, Lab-Control (*z*-ratio = 1.575, p-value = 0.1152) and 1-Day (*z*-ratio = 0.980, p-value = 0.3270) bees did not fan differently in either group composition type (Figure 3c). Although there was no significant interaction between individual treatment and group context on max bout fanning proportion (X^2^ = 2.185, p = 0.3354), we found a significant effect of both predictor variables as main effects (individual treatment, X^2^ = 45.050, p < 0.001; group composition type, X^2^ = 10.248, p = 0.0014).

### Individual bees move and interact differently depending on group context

Group composition significantly altered group movement (Figure 4a, X^2^ = 226.03, p < 0.001). Uniform 5-Day and mixed Lab-Control/5-Day groups had the lowest average group velocities (Figure 4a, mean group velocity ± standard deviation; 226.41 ± 61.19 and 237.43 ± 48.89, respectively). Mixed lab-Control/1-Day and 1-Day/5-Day groups had the highest average group velocities (Figure 4a, 357.82 ± 100.39 and 386.77 ± 129.59, respectively). When comparing individual velocity depending on mixed group partners, we found that individual Lab-Control bees had lower average velocity per second when they were grouped with 5-Day bees compared to when they were with 1-Day bees (Figure 4b, X^2^ = 38.447, p < 0.001). In contrast, the average velocity of individual 1-Day bees was not affected by the presence of mixed partners, as they moved at similar velocities regardless of partner identity (Figure 4c, X^2^= 0.10404, p = 0.75). Individual 5-Day bees also had lower average velocity per second when they were grouped with Lab-Control bees compared when they were grouped together with 1-Day bees (Figure 4d, X^2^ = 92.372, p < 0.001).

**Figure 4.**
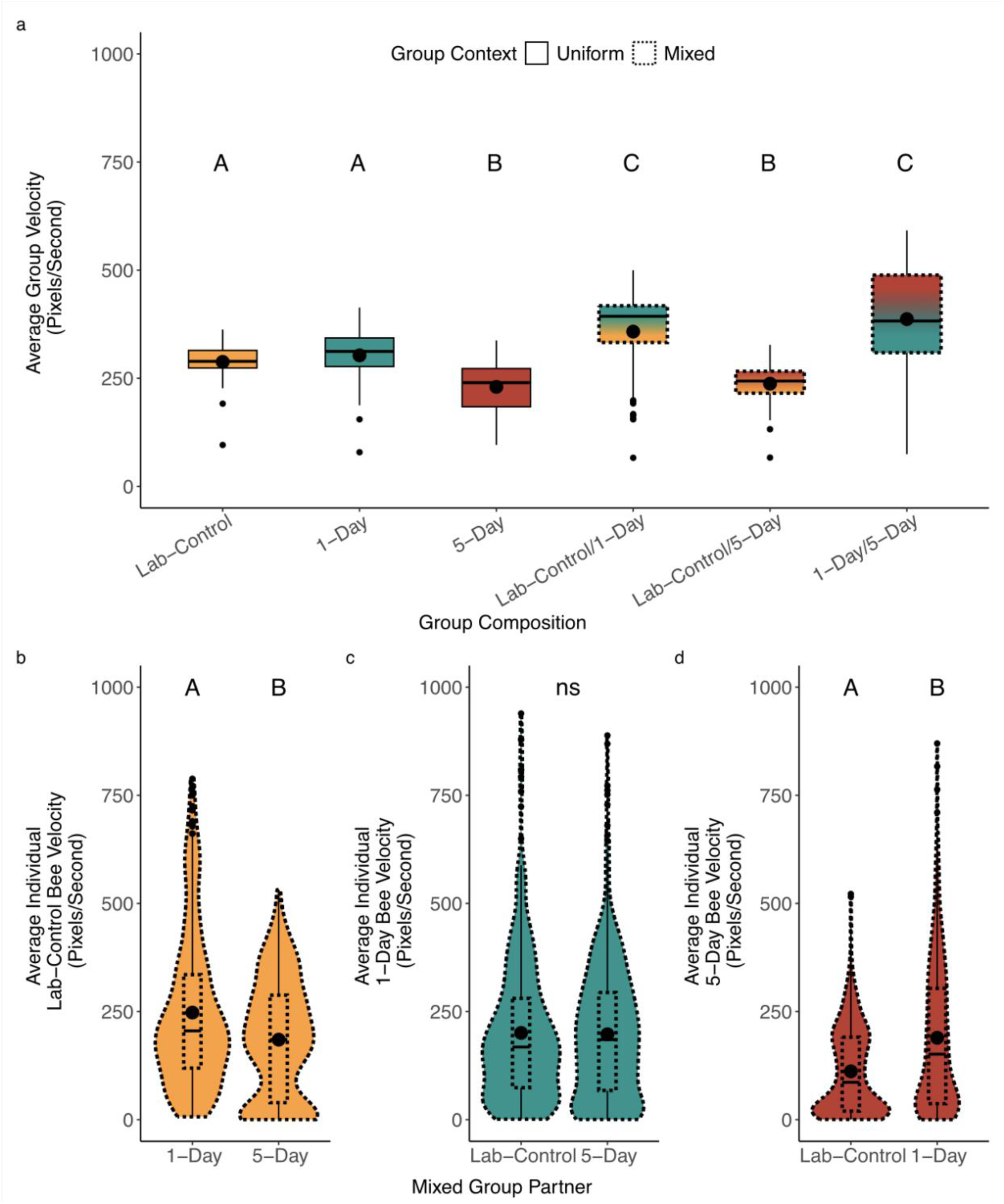
Individuals change their average movement velocity based on mixed group partner identity. N_Lab-Control_= 4, N_1-Day_ = 5, N_5-Day_ = 4, N_Lab-Control/5-Day_ = 3, N_Lab-Control/1-Day_ = 3, N_1-Day/5-Day_ = 5, for N = videos of 10 bees 4 minutes before the first fanning bout and 1 minute after for a total of 5 minutes. This meant that we had n = 95 individual bees per treatment group across all videos. a) Average group velocity across group composition. Lab-Control and 1-Day, 5-Day uniform and Lab-Control/5-Day, and Lab-Control/1Day and 1-Day/5-Day groups moved at similar group velocities to each other. GLM with a gamma distribution, X^2^ = 226.03, p < 0.0001; Tukey’s post hoc test, p < 0.05 b) Average Lab-Control bee velocity in the presence of 1-Day or 5-Day bees. Lab-Control bees moved significantly slower in the presence of 5-Day bees but not 1-Day bees. GLM with a gamma distribution, X^2^ = 38.447, p < 0.001 c) Average 1-Day bee velocity in the presence of lab-control or 5-Day bees. Individual 1-Day bee velocity was not impacted by the presence of lab-control or 5-Day bees. GLM with a gamma distribution, X^2^ = 0.10404, p = 0.747. d) Average 5-Day velocity in the presence of lab-control or 1-Day bees. 5-Day bees significantly move slower in the presence of lab-control bees. GLM with a gamma distribution, X^2^ = 92.372, p < 0.001. Letters indicate statistical significance. Center lines indicate median, while center dots indicate mean.

We found that group composition had a significant effect on the total number of head-to-head interactions per bee (Figure 5a, X^2^ = 50.23, p < 0.0001). Particularly, unform Lab-Control, uniform 5-Day, and mixed Lab-Control/1-Day groups had similar number of interactions to each other (Figure 5a, Tukey’s post hoc test, p < 0.05), while uniform 1-Day, mixed Lab-Control/5-Day, and mixed 1-Day/5-Day groups had similar number of interactions to each other (Figure 5a, Tukey’s post hoc test, p < 0.05). In mixed Lab-Control/5-Day and 1-Day/5-Day groups, interaction pairs dominated the proportion of interaction pairs we saw during videos, as we expected based on calculated combinatorics (Figure 5b, Tukey’s post hoc test, p < 0.001). However, in mixed Lab-Control/1-Day groups, mixed interaction pairs made up significantly fewer of all interactions than expected (Figure 5b, Chi-square test, X^2^ = 17.6, p < 0.0001).

**Figure 5.**
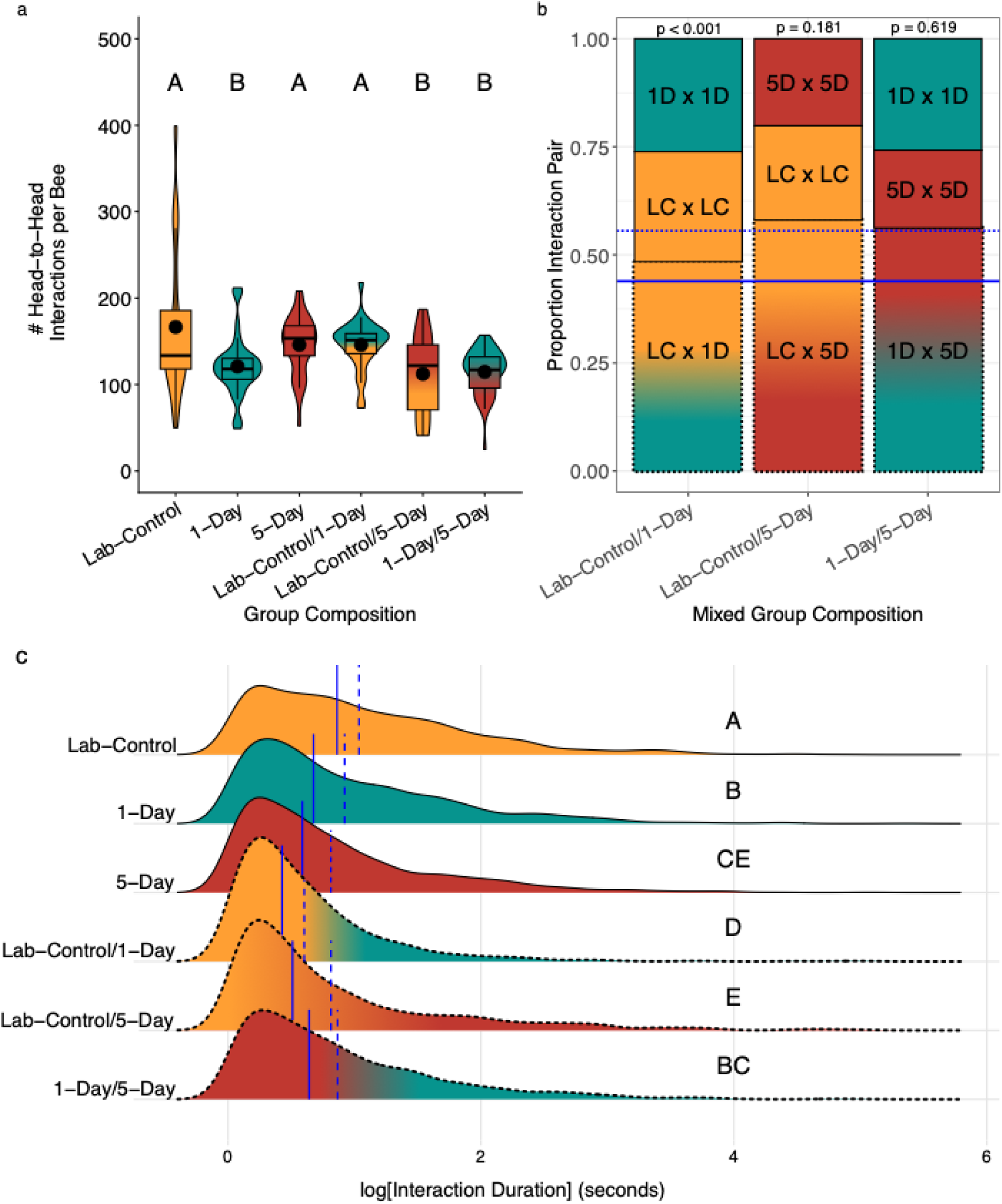
Group composition influences interaction dynamics across social contexts. N_Lab-Control_ = 4, N_1-Day_ = 5, N_5-Day_ = 4, N_Lab-Control/5-Day_ = 3, N_Lab-Control/1-Day_ = 3, N_1-Day/5-Day_ = 5, for N = videos of 10 bees 4 minutes before the first instance of fanning and 1 minute after for a total of 5 minutes a) Number of total head-to-head interactions per bee. Group composition had a significant effect, where Lab-Control, 5-Day, and Lab-Control/1-Day groups had similar number of interactions to each other and 1-Day, Lab-Control/5-Day, and 1-Day/5-Day groups had similar number of interactions to each other. GLM, X^2^=50.23, p <0.0001; Tukey’s post-hoc test, p < 0.05. Center lines indicate median, while center point indicate means b) Observed vs. expected interaction proportions in mixed groups based on combinatorial probabilities. Solid blue line indicates expected proportion for within-group interaction types combined, while dotted blue line indicates mixed group interaction types expected proportions based on combinatorial probabilities. Only the Lab-Control/1-Day group deviated significantly from what we expect combinatorially (Chi-Square test, χ² = 17.6, p < 0.001), with mixed pair interactions occurring less frequently than predicted c) Duration of head-to-head interactions across group composition. Lab-control and 1-Day/5-Day groups had the longest interaction duration, while 5-Day bees had the shortest. X-axis is log transformed to better visualize data spread. Dashed red line indicates the mean of interaction duration per group composition. Kruskal-Wallis test, X^2^ = 244.56, p < 0.001, Dunn’s Test p < 0.05. Letters indicate statistical significance. Solid, vertical blue line indicates median duration, while dotted, vertical blue line indicates mean duration.

Additionally, we found that group composition also significantly influenced the average duration of head-to-head interactions bees had within groups (Figure 5c, Kruskal-Wallis test, X^2^ = 244.56, p < 0.001; Dunn’s Test, p < 0.05). In uniform groups, lab-control bees had the longest lasting interactions (median, 2.37 seconds; mean, 4.48 ± 7.78 seconds) while 5-Day bees had the shortest (median, 1.80 seconds, 3.50 ± 6.94 seconds). In the mixed groups, lab-control/5-Day bees had the longest interaction duration (median, 1.67 seconds; mean 4.26 ± 10.91 seconds), followed by 1-Day/5-Day bees (median, 1.90 seconds; 3.75 seconds ± 7.59), and lab-control/1-Day bees had the shortest duration (median, 1.53 seconds; 2.62 ± 7.48).

### Individuals adjust their accessibility within social networks depending on group context

When comparing centrality across networks, we found no difference between all groups in betweenness (Figure 6a; LM, F = 0.5832, p = 0.7129). In line with what we saw in total number of head-to-head interactions, we found that uniform lab control groups had the highest network strength because we weighted strength by number of interactions per bee (Figure 6c; F = 5.9734, p < 0.0001; post-hoc comparison, p < 0.05). Furthermore, we found that lab control bees also had the highest network closeness, with most individuals occupying the center of the network, compared to all other groups (Figure 6b; 8.4103, p < 0.0001; post-hoc comparison, p < 0.05). Broadly, as groups became more disrupted, the centrality and connectedness of those groups significantly decreased.

**Figure 6.**
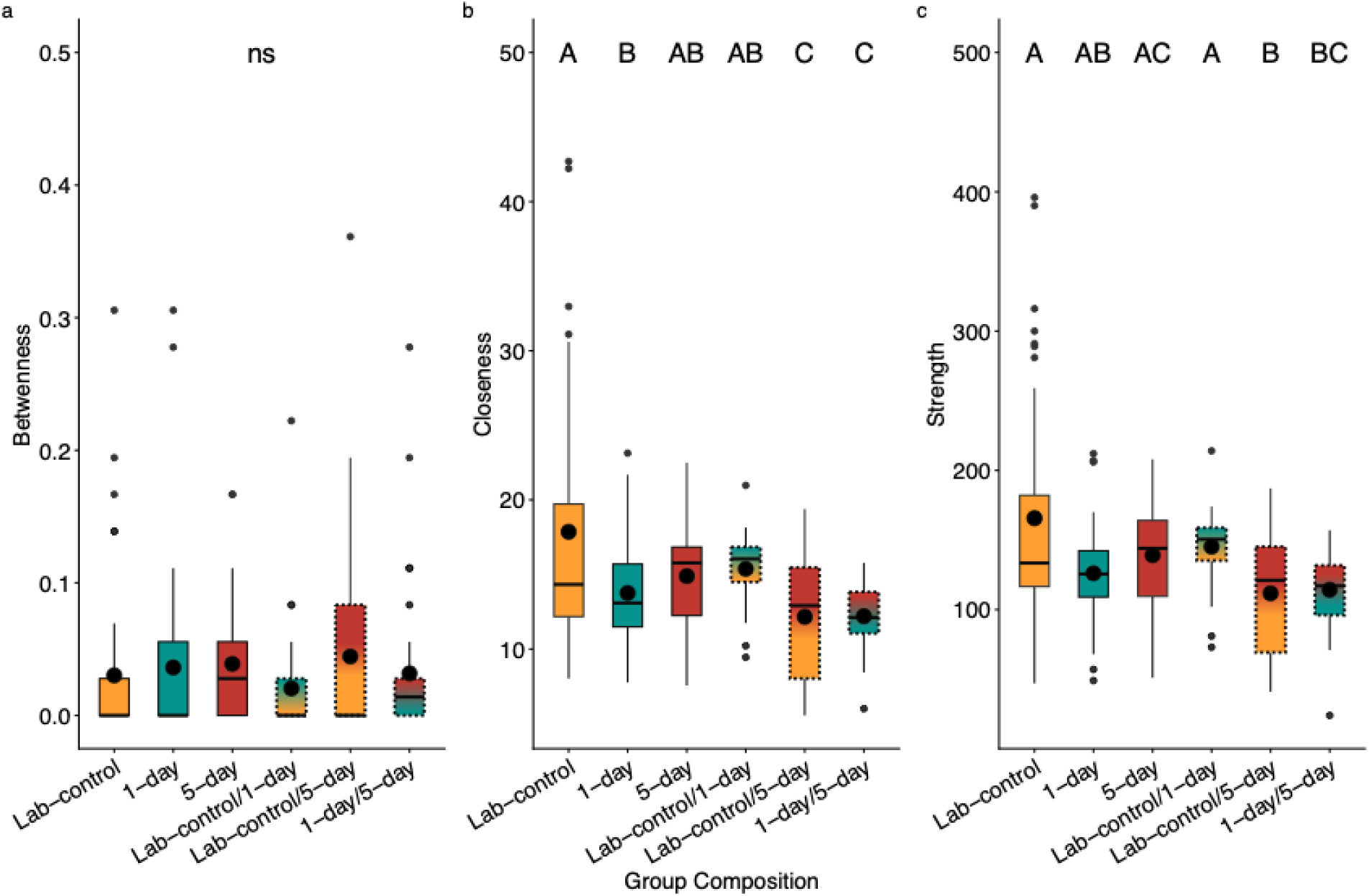
**Network closeness and strength vary depending on group composition**. N_lab control_ = 4, N_1-Day_ = 5, N_5-Day_ = 4, N_Lab-control/5-Day_ = 3, N_Lab-control/1-Day_ = 3, N_1-Day/5-Day_ = 5, for N = social networks constructed from ABCTracker data from videos of 10 bees 4 minutes before the first instance of fanning and 1 minute after for a total of 5 minutes a) we found no significant difference in betweenness across group composition. LM, F = 0.5832, p = 0.7129 b) Uniform lab-control groups had the highest closeness centrality of all groups. LM, F = 8.4103, p < 0.0001, Holm’s adjusted post-hoc comparison, p < 0.05 c) Uniform lab-control groups had the highest strength centrality of all groups. LM, F = 5.9734, p < 0.0001. Letters indicate statistical significance, where groups that share letters are not statistically different from one another. Center black dot in boxplots indicate mean. ns=not significant.

When we examined individual centrality across group contexts, we found that 5-Day bees decreased their network closeness and strength when in presence of other treatment bees (Figure 7e, F = 9.4379, p < 0.0001; pairwise comparison, p < 0.05; Figure 7e, F = 6.9032, p = 0.00188), moving away from the center of the social network. On the other hand, Lab-control and 1-Day bees did not alter their network strength when in the presence of other treatment groups (Figure 7b & 7d). 1-Day bees had significantly lower closeness when in the presence of 5-Day bees compared to when they were in the presence of Lab-Control bees (Figure 7e, F = 3.5312, p = 0.0341; pairwise comparison, p = 0.0341).

**Figure 7.**
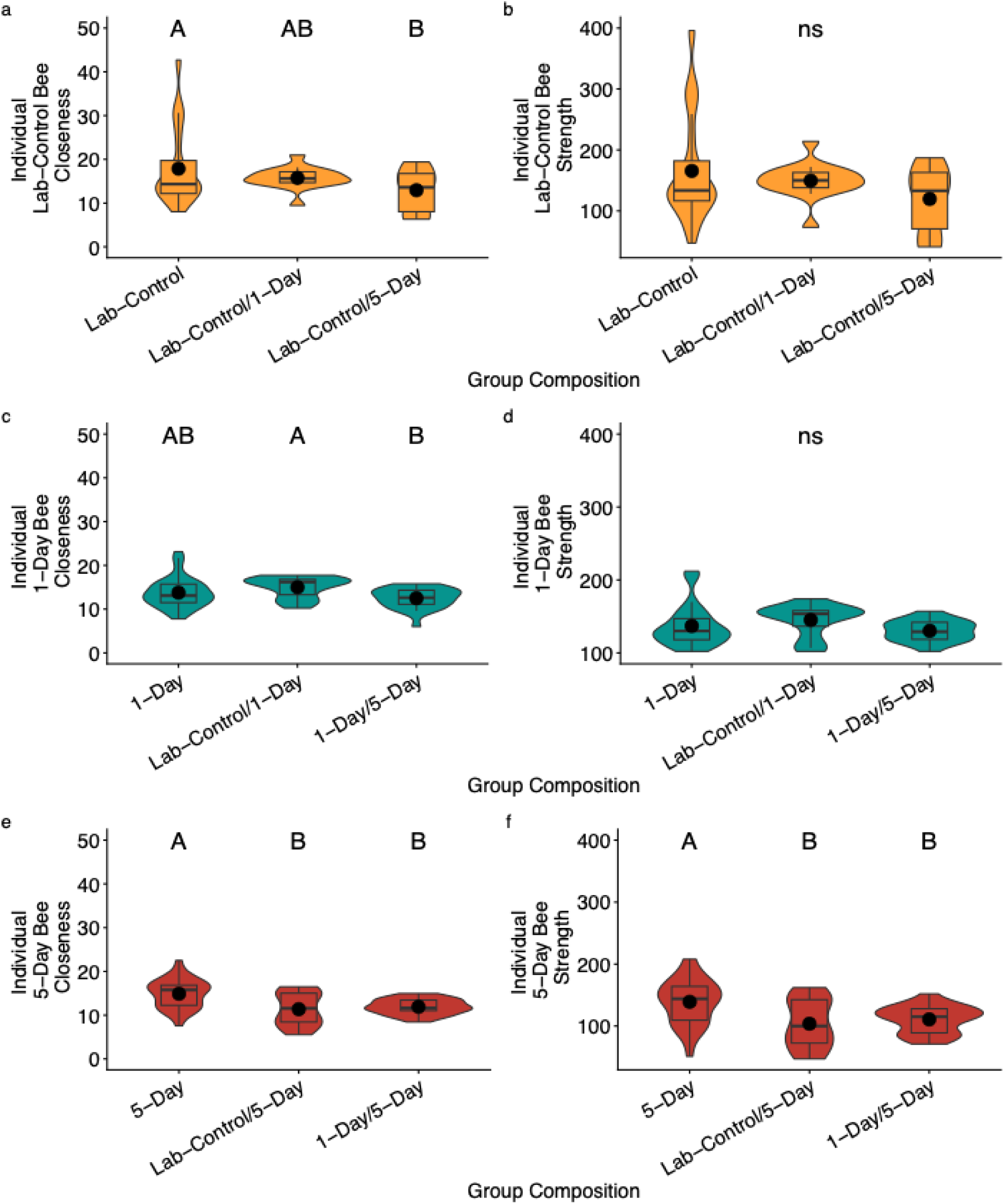
Individual bees adjust their social position depending on group context. N_lab control_ = 4, N_1-Day_ = 5, N_5-Day_ = 4, N_Lab-control/5-Day_ = 3, N_Lab-control/1-Day_ = 3, N_1-Day/5-Day_ = 5, for N = social networks constructed from ABCTracker data from videos of 10 bees 4 minutes before the first instance of fanning and 1 minute after for a total of 5 minutes. Y-axis scales differ among treatment groups because centrality metrics are calculated independently within each social network and are sensitive to network structure. Comparisons are therefore made within metrics across treatments rather than by absolute magnitude across networks a) Lab control bees had lower closeness scores when in the presence of 5-Day treatment bees. LM, F = 3.4064, p = 0.039, post-hoc comparison, p = 0.0357 b) No significant difference in lab control bee strength scores across group composition. LM, F = 1.2923, p =0.055 c) 1-Day bees had significantly lower closeness scores when paired with 5-Day bees compared to when they were paired with Lab-Control bees. LM, F = 0.3412, p = 0.0341; post hoc comparisons, p = 0.0305 d) No significant difference in 1-Day bees strength scores across group composition. LM, F= 2.061, p = 0.1343 e) 5-Day bees had significantly lower closeness scores when paired with other treatment bees. LM, F=1.0818, p = 0.00024; post hoc comparisons, p < 0.05 f) 5-Day bees had significantly lower strength scores when paired with other treatment bees. LM, F = 1.2413, p = 0.0012; post hoc-comparisons, p < 0.05. Center line indicates median, while center dots indicates mean. Letters indicate statistical significance where groups that do not share a letter are statistically different from one another.

Lab-Control bees also had significantly lowered closeness scores when paired with 5-Day bees compared to when they were with other Lab-Control bees (Figure 7a, F = 3.4064, p = 0.039; pairwise comparison, p < 0.05). There was no difference in individual betweenness across treatment groups (Supplementary Figure 2).

## Discussion

In this study, we found that honeybees can coordinate a fanning response with half of their group members impaired by shifting social organization. First, the likelihood of fanning in individual bees is significantly influenced by group context: 5-Day bee fanning performance can be rescued by the presence of Lab-Control bees in mixed group contexts (Figure 3). To explore some potential social mechanisms behind these observed rescue effects, we analyzed the network dynamics of our experimental groups at the group level and the individual level. Notably, we found that individual-level adjustments, particularly in movement velocity and interaction dynamics (Figure 4 & Figure 5), was correlated with this rescue effect. Individuals within mixed groups may have prioritized who they interacted with (Figure 5), how long they interacted with them (Figure 4), and how fast they moved depending on their mixed group partners (Figure 4). These results suggest that honeybees may be able to distinguish between disrupted and non-disrupted individuals and change their behavior to compensate for the disruption.

Group context also altered social networks and individual positions within those networks (Figure 6 & 7). Namely, 5-Day bees moved away from center social positions when in the presence of Lab-control or 1-Day treatment bees (Figure 7), potentially relinquishing leadership roles to healthier, better suited individuals. However, these 5-Day individual level adjustments were not sufficient to rescue fanning in 1-Day/5-Day groups. As a result, there may be a tipping point at which group function can no longer be rescued via individual adjustments in social position. Altogether, these findings lend support for our hypothesis that animal groups can be resilient to disruption, individual-level adjustments in social network positions appear to preserve group function, and after some threshold is met, such rescue effects can no longer sustain group function.

Behavioral plasticity is inextricably linked to the resilience of organisms, populations, and even ecosystems (Ghalambor et al., 2010; Júnior et al., 2022; Wong & Candolin, 2015). The ability to detect and respond to variations in the environment, such as group status, thermal change, or resource availability, is essential to buffer fitness costs (Júnior et al., 2022; Wong & Candolin, 2015). Social animals are particularly well poised to adjust to change because of decentralized ways of gathering and sharing information across group members that aid in consensus decision making (Bicca-Marques & Garber, 2005; Conradt & Roper, 2005; Couzin, 2009; King & Cowlishaw, 2007). For honeybees and other social insects, such flexibility is critical at the colony level when coordinating tasks with other workers (Garrison et al., 2018; Jeanson, 2019; Jeanson & Weidenmüller, 2014; Pinter-Wollman et al., 2011, 2013). Through interindividual behavioral plasticity, colonies can weather the fitness costs of disruptions, enabling task completion and colony homeostasis even under suboptimal conditions (Garrison et al., 2018; Jeanson & Weidenmüller, 2014; Neumann & Pinter-Wollman, 2019). For example, carpenter ants (*Camponotus pennsylvanicus*) keep their interaction networks stable by having a few discrete facilitators rapidly shuttle information between spatially distant clusters of workers (Modlmeier et al., 2019). Harvester ants (*Pogonomyrmex barbatus*) will adjust their foraging intensity depending on the rate at which other foragers return to the colony (Pinter-Wollman et al., 2013) and their volatile signals that indicate environmental temperature and humidity (Pagliara et al., 2018). The innate social structures of a social insect colony environment may therefore act as a safeguard when sections of the colony workforce are compromised by a variety of environmental stressors (Easton-Calabria et al., 2023; Garrison et al., 2018; Gordon, 2016; Naug, 2009).

Several physiological mechanisms could explain the results we see in our study. Social buffering is well-documented in vertebrates (Culbert et al., 2019; Edgar et al., 2015; Kiyokawa et al., 2007) and also occurs in social insects. For example, *Reticulitermes flavipes* termite workers paired with nestmate soldiers were protected from negative effects of competitor *R. virginicus* cues, such as increased mortality and delayed development. (Tian et al., 2017). In our study, it is possible that the presence of Lab-Control bees in mixed groups rescued 5-Day bees from the harmful effects of their oxytetracycline treatment, while the presence of 1-Day bees was not enough to convey this rescue effect. Another possible explanation for our findings could be disruption of chemical cues that mediate communication during interactions. Honeybee cuticular hydrocarbons (CHC) profiles are colony specific and are altered by antibiotic treatment, driving nestmate recognition (Vernier et al., 2020). Whether individual workers have unique, distinguishable CHC profiles is still heavily debated, although there is some evidence that honeybees can behaviorally respond to, discriminate, and learn individual CHCs (Châline et al., 2005; Dani et al., 2005). As a result, it is possible that in our study, Lab-Control bees could sense that 5-Day bees were disrupted and changed their behavior accordingly. Understanding whether and how honeybees detect each other’s physiological status and how that information influences individual decision-making and behavior will be important in unraveling the mechanisms behind group coordination.

Social network analysis (SNA) has been widely used to characterize collective behavior (Hasenjager & Dugatkin, 2015; Makagon et al., 2012; Omran & and van Etten, 2007; Pinheiro et al., 2016; Wasserman & Faust, 1994; Wey et al., 2008). Thus far, SNA studies have classified the social organization in a wide variety of taxa, including troops of primates (Kasper & Voelkl, 2009; King & Sueur, 2011; Sueur et al., 2010), flocks of birds (Curry & Dunbar, 2013; Farine et al., 2012; Ren et al., 2018), schools of fish (Jolles et al., 2017; Katz et al., 2011), and colonies and swarms of insects (Alwash & Levine, 2019; Buhl et al., 2011; Naug, 2009; Pinter-Wollman et al., 2013; Stroeymeyt et al., 2018; Topaz et al., 2012; Waters & Fewell, 2012). The continual advancement in SNA technologies have provided exciting avenues to ask more nuanced questions about the resiliency of animal networks and the role that individuals play in that resiliency, especially within the contexts of rapid and compounding environmental change (Buchholz et al., 2019; Komdeur & Ma, 2021; Reed et al., 2024).

As social network analysis technologies continue to advance and become more accessible to researchers, we have powerful tools at our disposal to disentangle the fine-scale social dynamics that underlie collective behavior (Pereira et al., 2022; Smith & Pinter-Wollman, 2021) and how human-driven environmental change could impact these dynamics (Buchholz et al., 2019; Komdeur & Ma, 2021; Michelangeli et al., 2022). Applying these tools to study insect societies enables a deeper understanding of how colonies maintain functionality under stress, revealing individual-level plasticity that can sustain group-level resilience. Colonies undoubtedly have a tipping point at which workers can no longer maintain homeostasis. In the face for rapid, human driven environmental change, it is imperative to examine how and when collective function is preserved and breaks down. Future interdisciplinary work that integrates behavior, physiology, and ecology will be crucial in uncovering the full suite of mechanisms that support coordination under disruption. By continuing to study social insects, we can better understand plasticity and resilience in social animals to better predict the futures of social animal populations.

## Acknowledgements

We would like to thank Natalia Nowakowski for her assistance in collecting data. Thank you to the members of the Cook Lab as well as to Dr. Noa Pinter-Wollman for helpful feedback. This work was supported by National Science Foundation Integrative Organismal Systems grant (#2212640) awarded to C.N.C. and U.S. Department of Agriculture North Central Sustainable Agriculture Research and Education grant (GNC23-370) awarded to J.B.N. Data and code are provided on FigShare.

## Author Contribution

Conceptualization, J.B.N and C.N.C.; Methodology, J.B.N, C.E.L., C.N.C.; Validation, J.B.N. and C.E.L.; Formal Analysis, J.B.N and C.E.L.; Investigation, J.B.N.; Resources, C.N.C.; Data curation, J.B.N. and C.E.L; Writing – Original Draft, J.B.N and C.N.C.; Writing – Review & Editing, J.B.N , C.N.C. and C.E.L.; Visualization, J.B.N and C.N.C.; Supervision, C.N.C.; Project Administration, C.N.C.

## Supplemental Information

Document 1.

## References

Abram, P. K., Boivin, G., Moiroux, J., & Brodeur, J. (2017). Behavioural effects of temperature on ectothermic animals: Unifying thermal physiology and behavioural plasticity. Biological Reviews, 92(4), 1859–1876. 10.1111/brv.12312

Alwash, N., & Levine, J. D. (2019). Network analyses reveal structure in insect social groups. Current Opinion in Insect Science, Global Change Biology • Molecular Physiology, 35, 54–59. 10.1016/j.cois.2019.07.001

Bentzur, A., Ben-Shaanan, S., Benichou, J. I. C., Costi, E., Levi, M., Ilany, A., & Shohat-Ophir, G. (2021). Early Life Experience Shapes Male Behavior and Social Networks in Drosophila. Current Biology, 31(3), 486–501.e3. 10.1016/j.cub.2020.10.060

Beshers, S. N., Huang, Z. Y., Oono, Y., & Robinson, G. E. (2001). Social Inhibition and the Regulation of Temporal Polyethism in Honey Bees. Journal of Theoretical Biology, 213(3), 461–479. 10.1006/jtbi.2001.2427

Bicca-Marques, J. C., & Garber, P. A. (2005). Use of Social and Ecological Information in Tamarin Foraging Decisions. International Journal of Primatology, 26(6), 1321–1344. 10.1007/s10764-005-8855-9

Blumstein, D. T., Hayes, L. D., & Pinter-Wollman, N. (2023). Social consequences of rapid environmental change. *Trends in Ecology & Evolution*, Special Issue: Animal Behaviour in a Changing World, 38(4), 337–345. 10.1016/j.tree.2022.11.005

Bradshaw, W. E., & Holzapfel, C. M. (2010). Light, Time, and the Physiology of Biotic Response to Rapid Climate Change in Animals. Annual Review of Physiology, 72(Volume 72, 2010), 147–166. 10.1146/annurev-physiol-021909-135837

Buchholz, R., Banusiewicz, J. D., Burgess, S., Crocker-Buta, S., Eveland, L., & Fuller, L. (2019). Behavioural research priorities for the study of animal response to climate change. Animal Behaviour, 150, 127–137. 10.1016/j.anbehav.2019.02.005

Buhl, C., Sword, G. A., Clissold, F. J., & Simpson, S. J. (2011). Group structure in locust migratory bands. *Behavioral Ecology and Sociobiology*, 65(2), 265–273. 10.1007/s00265-010-1041-x

Cannon, W. B. (1929). Organization for physiological homeostasis. *Physiological Reviews*, S(3), 399–431. 10.1152/physrev.1929.9.3.399

Châline, N., Sandoz, J.-C., Martin, S. J., Ratnieks, F. L. W., & Jones, G. R. (2005). Learning and Discrimination of Individual Cuticular Hydrocarbons by Honeybees (Apis mellifera). Chemical Senses, 30(4), 327–335. 10.1093/chemse/bji027

Conradt, L., & Roper, T. J. (2005). Consensus decision making in animals. Trends in Ecology & Evolution, 20(8), 449–456. 10.1016/j.tree.2005.05.008

Cook, C. N., & Breed, M. D. (2013). Social context influences the initiation and threshold of thermoregulatory behaviour in honeybees. Animal Behaviour, 86(2), 323–329. 10.1016/j.anbehav.2013.05.021

Cook, C. N., Kaspar, R. E., Flaxman, S. M., & Breed, M. D. (2016). Rapidly changing environment modulates the thermoregulatory fanning response in honeybee groups. Animal Behaviour, 115, 237–243. 10.1016/j.anbehav.2016.03.014

Cook, C. N., Lemanski, N. J., Mosqueiro, T., Ozturk, C., Gadau, J., Pinter-Wollman, N., & Smith, B. H. (2020). Individual learning phenotypes drive collective behavior. Proceedings of the National Academy of Sciences, 117(30), 17949. 10.1073/pnas.1920554117

Couzin, I. D. (2009). Collective cognition in animal groups. Trends in Cognitive Sciences, 13(1), 36–43. 10.1016/j.tics.2008.10.002

Csardi, G., & Nepusz, T. (2006). The igraph software package for complex network research. InterJournal, Complex Systems. https://igraph.org

Csárdi, G., Nepusz, T., Müller, K., Horvát, S., Traag, V., Zanini, F., & Noom, D. (2025). *igraph for R: R interface of the igraph library for graph theory and network analysis* [Computer software]. Zenodo. 10.5281/zenodo.14736815

Culbert, B. M., Gilmour, K. M., & Balshine, S. (2019). Social buffering of stress in a group-living fish. Proceedings of the Royal Society B: Biological Sciences, 286(1910), 20191626. 10.1098/rspb.2019.1626

Curry, O., & Dunbar, R. I. M. (2013). Do Birds of a Feather Flock Together? Human Nature, 24(3), 336–347. 10.1007/s12110-013-9174-z

Dani, F. R., Jones, G. R., Corsi, S., Beard, R., Pradella, D., & Turillazzi, S. (2005). Nestmate Recognition Cues in the Honey Bee: Differential Importance of Cuticular Alkanes and Alkenes. Chemical Senses, 30(6), 477–489. 10.1093/chemse/bji040

Davidson, J. D., Vishwakarma, M., & Smith, M. L. (2021). Hierarchical Approach for Comparing Collective Behavior Across Scales: Cellular Systems to Honey Bee Colonies. *Frontiers in Ecology and Evolution*, S. 10.3389/fevo.2021.581222

Davies, K. J. A. (2016). Adaptive homeostasis. *Molecular Aspects of Medicine*, Hormetic and Regulatory Effects of Lipid Oxidation Products, *49*, 1–7. 10.1016/j.mam.2016.04.007

del Mar Delgado, M., Miranda, M., Alvarez, S. J., Gurarie, E., Fagan, W. F., Penteriani, V., di Virgilio, A., & Morales, J. M. (2018). The importance of individual variation in the dynamics of animal collective movements. Philosophical Transactions of the Royal Society B: Biological Sciences, 373(1746), 20170008. 10.1098/rstb.2017.0008

Easton-Calabria, A. C., Thuma, J. A., Cronin, K., Melone, G., Laskowski, M., Smith, M. A. Y., Pasadyn, C. L., de Bivort, B. L., & Crall, J. D. (2023). Colony size buffers interactions between neonicotinoid exposure and cold stress in bumblebees. Proceedings of the Royal Society B: Biological Sciences, 290(2003), 20230555. 10.1098/rspb.2023.0555

Edgar, J., Held, S., Paul, E., Pettersson, I., I’Anson Price, R., & Nicol, C. (2015). Social buffering in a bird. Animal Behaviour, 105, 11–19. 10.1016/j.anbehav.2015.04.007

Egley, R. L., & Breed, M. D. (2013). The Fanner Honey Bee: Behavioral Variability and Environmental Cues in Workers Performing a Specialized Task. Journal of Insect Behavior, 2c(2), 238–245. 10.1007/s10905-012-9357-1

Farine, D. R., Garroway, C. J., & Sheldon, B. C. (2012). Social network analysis of mixed-species flocks: Exploring the structure and evolution of interspecific social behaviour. Animal Behaviour, 84(5), 1271–1277. 10.1016/j.anbehav.2012.08.008

Farine, D. R., & Whitehead, H. (2015). Constructing, conducting and interpreting animal social network analysis. The Journal of Animal Ecology, 84(5), 1144–1163. 10.1111/1365-2656.12418

Fernandez, M. S. A., Vignal, C., & Soula, H. A. (2017). Impact of group size and social composition on group vocal activity and acoustic network in a social songbird. Animal Behaviour, 127, 163–178. 10.1016/j.anbehav.2017.03.013

Garnier, S., Murphy, T., Lutz, M., Hurme, E., Leblanc, S., & Couzin, I. D. (2013). Stability and Responsiveness in a Self-Organized Living Architecture. *PLOS Computational Biology*, S(3), e1002984. 10.1371/journal.pcbi.1002984

Garrison, L. K., Kleineidam, C. J., & Weidenmüller, A. (2018). Behavioral flexibility promotes collective consistency in a social insect. Scientific Reports, 8(1), 15836. 10.1038/s41598-018-33917-7

Ghalambor, C. K., Angeloni, L. M., & Carroll, S. P. (2010). Behavior as Phenotypic Plasticity. Evolutionary Behavioral Ecology, 90–107.

Gordon, D. M. (2016). From division of labor to the collective behavior of social insects. Behavioral Ecology and Sociobiology, 70(7), 1101–1108. 10.1007/s00265-015-2045-3

Gordon, D. M. (2023). Collective behavior in relation with changing environments: Dynamics, modularity, and agency. Evolution & Development, 25(6), 430–438. 10.1111/ede.12439

Grüter, C., & Farina, W. M. (2009). The honeybee waggle dance: Can we follow the steps? Trends in Ecology & Evolution, 24(5), 242–247. 10.1016/j.tree.2008.12.007

Hasenjager, M. J., & Dugatkin, L. A. (2015). Chapter Three—Social Network Analysis in Behavioral Ecology. In M. Naguib, H. J. Brockmann, J. C. Mitani, L. W. Simmons, L. Barrett, S. Healy, & P. J. B. Slater (Eds.), Advances in the Study of Behavior (Vol. 47, pp. 39–114). Academic Press. 10.1016/bs.asb.2015.02.003

Herbert-Read, J. E., Krause, S., Morrell, L. J., Schaerf, T. M., Krause, J., & Ward, A. J. W. (2013). The role of individuality in collective group movement. Proceedings of the Royal Society B: Biological Sciences, 280(1752), 20122564. 10.1098/rspb.2012.2564

Hoogland, J. L. (1983). Nepotism and alarm calling in the black-tailed prairie dog (Cynomys ludovicianus). Animal Behaviour, 31(2), 472–479. 10.1016/S0003-3472(83)80068-2

Hoogland, J. L. (1995). The Black-Tailed Prairie Dog: Social Life of a Burrowing Mammal. University of Chicago Press.

Huang, Z.-Y., & Robinson, G. E. (1992). Honeybee colony integration: Worker-worker interactions mediate hormonally regulated plasticity in division of labor. Proceedings of the National Academy of Sciences, 89(24), 11726–11729. 10.1073/pnas.89.24.11726

Huang, Z.-Y., & Robinson, G. E. (1996). Regulation of honey bee division of labor by colony age demography. Behavioral Ecology and Sociobiology, 39(3), 147–158. 10.1007/s002650050276

Jäger, H. Y., Han, C. S., & Dingemanse, N. J. (2019). Social experiences shape behavioral individuality and within-individual stability. Behavioral Ecology, 30(4), 1012–1019. 10.1093/beheco/arz042

Jeanson, R. (2019). Within-individual behavioural variability and division of labour in social insects. Journal of Experimental Biology, 222(10), jeb190868. 10.1242/jeb.190868

Jeanson, R., & Weidenmüller, A. (2014). Interindividual variability in social insects – proximate causes and ultimate consequences. Biological Reviews, 89(3), 671–687. 10.1111/brv.12074

Jolles, J. W., Boogert, N. J., Sridhar, V. H., Couzin, I. D., & Manica, A. (2017). Consistent Individual Differences Drive Collective Behavior and Group Functioning of Schooling Fish. Current Biology, 27(18), 2862–2868.e7. 10.1016/j.cub.2017.08.004

Jolles, J. W., King, A. J., & Killen, S. S. (2020). The Role of Individual Heterogeneity in Collective Animal Behaviour. Trends in Ecology & Evolution, 35(3), 278–291. 10.1016/j.tree.2019.11.001

Júnior, E. C. B., Rios, V. P., Dodonov, P., Vilela, B., & Japyassú, H. F. (2022). Effect of behavioural plasticity and environmental properties on the resilience of communities under habitat loss and fragmentation. Ecological Modelling, 472, 110071. 10.1016/j.ecolmodel.2022.110071

Kaspar, R. E., Cook, C. N., & Breed, M. D. (2018). Experienced individuals influence the thermoregulatory fanning behaviour in honey bee colonies. Animal Behaviour, 142, 69–76. 10.1016/j.anbehav.2018.06.004

Kasper, C., & Voelkl, B. (2009). A social network analysis of primate groups. Primates, 50(4), 343–356. 10.1007/s10329-009-0153-2

Kassahn, K. S., Crozier, R. H., Pörtner, H. O., & Caley, M. J. (2009). Animal performance and stress: Responses and tolerance limits at different levels of biological organisation. Biological Reviews, 84(2), 277–292. 10.1111/j.1469-185X.2008.00073.x

Katz, Y., Tunstrøm, K., Ioannou, C. C., Huepe, C., & Couzin, I. D. (2011). Inferring the structure and dynamics of interactions in schooling fish. Proceedings of the National Academy of Sciences, 108(46), 18720–18725. 10.1073/pnas.1107583108

Kay, T., Liberti, J., Richardson, T. O., McKenzie, S. K., Weitekamp, C. A., La Mendola, C., Rüegg, M., Kesner, L., Szombathy, N., McGregor, S., Romiguier, J., Engel, P., & Keller, L. (2023). Social network position is a major predictor of ant behavior, microbiota composition, and brain gene expression. PLOS Biology, 21(7), e3002203. 10.1371/journal.pbio.3002203

Killen, S. S., Marras, S., Metcalfe, N. B., McKenzie, D. J., & Domenici, P. (2013). Environmental stressors alter relationships between physiology and behaviour. Trends in Ecology & Evolution, 28(11), 651–658. 10.1016/j.tree.2013.05.005

King, A. J., & Cowlishaw, G. (2007). When to use social information: The advantage of large group size in individual decision making. Biology Letters, 3(2), 137–139. 10.1098/rsbl.2007.0017

King, A. J., & Sueur, C. (2011). Where Next? Group Coordination and Collective Decision Making by Primates. International Journal of Primatology, 32(6), 1245–1267. 10.1007/s10764-011-9526-7

Kiyokawa, Y., Takeuchi, Y., & Mori, Y. (2007). Two types of social buffering differentially mitigate conditioned fear responses. European Journal of Neuroscience, 26(12), 3606–3613. 10.1111/j.1460-9568.2007.05969.x

Komdeur, J., & Ma, L. (2021). Keeping up with environmental change: The importance of sociality. Ethology, 127(10), 790–807. 10.1111/eth.13200

Koolhaas, J. M. (2008). Coping style and immunity in animals: Making sense of individual variation. *Brain, Behavior, and Immunity*, Personality and Disease, 22(5), 662–667. 10.1016/j.bbi.2007.11.006

Koolhaas, J. M., de Boer, S. F., Coppens, C. M., & Buwalda, B. (2010). Neuroendocrinology of coping styles: Towards understanding the biology of individual variation. Frontiers in Neuroendocrinology, 31(3), 307–321. 10.1016/j.yfrne.2010.04.001

Krause, J., Krause, P. of F. B. and E. J., Ruxton, G. D., Ruxton, G., & Ruxton, I. G. (2002). Living in Groups. OUP Oxford.

Lambert, C. E., Vazquez, K., Nelson, Z. P., & Cook, C. N. (2025). Physical Interactions Shape Collective Thermoregulatory Behavior in Honey Bees. *Behavioral Ecology*, araf106. 10.1093/beheco/araf106

Laub, E. C., Pinter-Wollman, N., & Tibbetts, E. A. (2024). Personality and body mass impact social group formation and function in paper wasps. Animal Behaviour, 213, 207–218. 10.1016/j.anbehav.2024.03.020

LeBoeuf, A. C., & Grozinger, C. M. (2014). Me and we: The interplay between individual and group behavioral variation in social collectives. Current Opinion in Insect Science, 5, 16–24. 10.1016/j.cois.2014.09.010

Lemanski, N. J., Cook, C. N., Smith, B. H., & Pinter-Wollman, N. (2019). A Multiscale Review of Behavioral Variation in Collective Foraging Behavior in Honey Bees. Insects, 10(11), Article 11. 10.3390/insects10110370

Li, Z., Bhat, B., Frank, E. T., Oliveira-Honorato, T., Azuma, F., Bachmann, V., Parker, D. J., Schmitt, T., Economo, E. P., & Ulrich, Y. (2023). Behavioural individuality determines infection risk in clonal ant colonies. Nature Communications, 14(1), 5233. 10.1038/s41467-023-40983-7

Lindauer, M. (1955). The Water Economy and Temperature Regulation of the Honeybee Colony. Bee World, 36(5), 81–92. 10.1080/0005772X.1955.11094876

Makagon, M. M., McCowan, B., & Mench, J. A. (2012). How can social network analysis contribute to social behavior research in applied ethology? *Applied Animal Behaviour Science*, Special Issue: Living In Large Groups, 138(3), 152–161. 10.1016/j.applanim.2012.02.003

Masood, F., Thebeau, J. M., Cloet, A., Kozii, I. V., Zabrodski, M. W., Biganski, S., Liang, J., Marta Guarna, M., Simko, E., Ruzzini, A., & Wood, S. C. (2022). Evaluating approved and alternative treatments against an oxytetracycline-resistant bacterium responsible for European foulbrood disease in honey bees. Scientific Reports, 12, 5906. 10.1038/s41598-022-09796-4

McHenry, L. C., Schürch, R., Johnson, L. E., Ohlinger, B. D., & Couvillon, M. J. (2025). Individuality impacts communication success in honey bees. Current Biology, 35(4), R137–R138. 10.1016/j.cub.2024.12.047

Michelangeli, M., Martin, J. M., Pinter-Wollman, N., Ioannou, C. C., McCallum, E. S., Bertram, M. G., & Brodin, T. (2022). Predicting the impacts of chemical pollutants on animal groups. Trends in Ecology & Evolution, 37(9), 789–802. 10.1016/j.tree.2022.05.009

Modlmeier, A. P., Colman, E., Hanks, E. M., Bringenberg, R., Bansal, S., & Hughes, D. P. (2019). Ant colonies maintain social homeostasis in the face of decreased density. eLife, 8, e38473. 10.7554/eLife.38473

Nardone, A., Ronchi, B., Lacetera, N., Ranieri, M. S., & Bernabucci, U. (2010). Effects of climate changes on animal production and sustainability of livestock systems. Livestock Science, 10th World Conference on Animal Production (WCAP), 130(1), 57–69. 10.1016/j.livsci.2010.02.011

Naug, D. (2009). Structure and resilience of the social network in an insect colony as a function of colony size. *Behavioral Ecology and Sociobiology*, 63(7), 1023–1028. 10.1007/s00265-009-0721-x

Neumann, K. M., & Pinter-Wollman, N. (2019). Collective responses to heterospecifics emerge from individual differences in aggression. Behavioral Ecology, 30(3), 801– 808. 10.1093/beheco/arz017

Nguyen, J. B., & Cook, C. N. (2025). Disruption of collective behaviour correlates with reduced interaction efficiency. Proceedings of the Royal Society B: Biological Sciences, 2S2. 10.1098/rspb.2025.0039

Oliveira, R. F. (2009). Social behavior in context: Hormonal modulation of behavioral plasticity and social competence. Integrative and Comparative Biology, *49*(4), 423–440. 10.1093/icb/icp055

Omran, El. E., C and van Etten, J. (2007). Spatial-Data Sharing: Applying Social-Network Analysis to study individual and collective behaviour. International Journal of Geographical Information Science, 21(6), 699–714. 10.1080/13658810601135726

Ono, M., Igarashi, T., Ohno, E., & Sasaki, M. (1995). Unusual thermal defence by a honeybee against mass attack by hornets. Nature, 377(6547), 334–336. 10.1038/377334a0

Ono, M., Okada, I., & Sasaki, M. (1987). Heat production by balling in the Japanese honeybee,Apis cerana japonica as a defensive behavior against the hornet,Vespa simillima xanthoptera (Hymenoptera: Vespidae). Experientia, 43(9), 1031–1034. 10.1007/BF01952231

Ortiz-Alvarado, Y., Clark, D. R., Vega-Melendez, C. J., Flores-Cruz, Z., Domingez-Bello, M. G., & Giray, T. (2020). Antibiotics in hives and their effects on honey bee physiology and behavioral development. *Biology Open*, 9(bio053884). 10.1242/bio.053884

Pagliara, R., Gordon, D. M., & Leonard, N. E. (2018). Regulation of harvester ant foraging as a closed-loop excitable system. PLOS Computational Biology, 14(12), e1006200. 10.1371/journal.pcbi.1006200

Pereira, T. D., Tabris, N., Matsliah, A., Turner, D. M., Li, J., Ravindranath, S., Papadoyannis, E. S., Normand, E., Deutsch, D. S., Wang, Z. Y., McKenzie-Smith, G. C., Mitelut, C. C., Castro, M. D., D’Uva, J., Kislin, M., Sanes, D. H., Kocher, S. D., Wang, S. S.-H., Falkner, A. L., … Murthy, M. (2022). SLEAP: A deep learning system for multi-animal pose tracking. Nature Methods, *19*(4), 486–495. 10.1038/s41592-022-01426-1

Pezzulo, G., Rigoli, F., & Friston, K. (2015). Active Inference, homeostatic regulation and adaptive behavioural control. Progress in Neurobiology, 134, 17–35. 10.1016/j.pneurobio.2015.09.001

Pinheiro, F. L., Santos, F. C., & Pacheco, J. M. (2016). Linking Individual and Collective Behavior in Adaptive Social Networks. Physical Review Letters, 116(12), 128702. 10.1103/PhysRevLett.116.128702

Pinter-Wollman, N., Bala, A., Merrell, A., Queirolo, J., Stumpe, M. C., Holmes, S., & Gordon, D. M. (2013). Harvester ants use interactions to regulate forager activation and availability. Animal Behaviour, *86*(1), 197–207. 10.1016/j.anbehav.2013.05.012

Pinter-Wollman, N., Wollman, R., Guetz, A., Holmes, S., & Gordon, D. M. (2011). The effect of individual variation on the structure and function of interaction networks in harvester ants. Journal of The Royal Society Interface, 8(64), 1562–1573. 10.1098/rsif.2011.0059

Radchuk, V., Reed, T., Teplitsky, C., van de Pol, M., Charmantier, A., Hassall, C., Adamík, P., Adriaensen, F., Ahola, M. P., Arcese, P., Miguel Avilés, J., Balbontin, J., Berg, K. S., Borras, A., Burthe, S., Clobert, J., Dehnhard, N., de Lope, F., Dhondt, A. A., … Kramer-Schadt, S. (2019). Adaptive responses of animals to climate change are most likely insufficient. Nature Communications, 10(1), 3109. 10.1038/s41467-019-10924-4

Raymann, K., Bobay, L.-M., & Moran, N. A. (2018). Antibiotics reduce genetic diversity of core species in the honeybee gut microbiome. Molecular Ecology, 27(8), 2057– 2066. 10.1111/mec.14434

Raymann, K., Shaffer, Z., & Moran, N. A. (2017). Antibiotic exposure perturbs the gut microbiota and elevates mortality in honeybees. PLoS Biology, 15(3). 10.1371/journal.pbio.2001861

Reed, J. M., Wolfe, B. E., & Romero, L. M. (2024). Is resilience a unifying concept for the biological sciences? iScience, 27(5), 109478. 10.1016/j.isci.2024.109478

Reid, C. R., Lutz, M. J., Powell, S., Kao, A. B., Couzin, I. D., & Garnier, S. (2015). Army ants dynamically adjust living bridges in response to a cost–benefit trade-off. Proceedings of the National Academy of Sciences, 112(49), 15113–15118. 10.1073/pnas.1512241112

Ren, J., Sun, W., Manocha, D., Li, A., & Jin, X. (2018). Stable information transfer network facilitates the emergence of collective behavior of bird flocks. *Physical Review E*, S8(5), 052309. 10.1103/PhysRevE.98.052309

Rice, L., Tate, S., Farynyk, D., Sun, J., Chism, G., Charbonneau, D., Fasciano, T., Dornhaus, A., & Shin, M. C. (2020). *ABCTracker: An easy-to-use, cloud-based application for tracking multiple objects* (arXiv:2001.10072). arXiv. 10.48550/arXiv.2001.10072

Richardson, T. O., Stroeymeyt, N., Crespi, A., & Keller, L. (2022). Two simple movement mechanisms for spatial division of labour in social insects. Nature Communications, 13(1), 6985. 10.1038/s41467-022-34706-7

Riley, J. R., Greggers, U., Smith, A. D., Reynolds, D. R., & Menzel, R. (2005). The flight paths of honeybees recruited by the waggle dance. Nature, 435(7039), 205–207.10.1038/nature03526

Seeley, T. D., & Buhrman, S. C. (1999). Group decision making in swarms of honey bees. Behavioral Ecology and Sociobiology, 45(1), 19–31. 10.1007/s002650050536

Seeley, T. D., Camazine, S., & Sneyd, J. (1991). Collective decision-making in honey bees: How colonies choose among nectar sources. Behavioral Ecology and Sociobiology, 28(4), 277–290. 10.1007/BF00175101

Seeley, T. D., & Kirk Visscher, P. (2004). Group decision making in nest-site selection by honey bees. Apidologie, 35(2), 101–116. 10.1051/apido:2004004

Smith, J. E., & Pinter-Wollman, N. (2021). Observing the unwatchable: Integrating automated sensing, naturalistic observations and animal social network analysis in the age of big data. *Journal of Animal Ecology*, 90(1), 62–75. 10.1111/1365-2656.13362

Stroeymeyt, N., Grasse, A. V., Crespi, A., Mersch, D. P., Cremer, S., & Keller, L. (2018). Social network plasticity decreases disease transmission in a eusocial insect. Science, 3c2(6417), 941–945. 10.1126/science.aat4793

Sueur, C., Deneubourg, J.-L., & Petit, O. (2010). Sequence of quorums during collective decision making in macaques. *Behavioral Ecology and Sociobiology*, 64(11), 1875– 1885. 10.1007/s00265-010-0999-8

Sugahara, M., Nishimura, Y., & Sakamoto, F. (2012). Differences in Heat Sensitivity between Japanese Honeybees and Hornets Under High Carbon Dioxide and Humidity Conditions Inside Bee Balls. Zoological Science, 29(1), 30–36. 10.2108/zsj.29.30

Swain, A., Williams, S. D., Di Felice, L. J., & Hobson, E. A. (2022). Interactions and information: Exploring task allocation in ant colonies using network analysis. Animal Behaviour, *18S*, 69–81. 10.1016/j.anbehav.2022.04.015

Thorogood, R., Mustonen, V., Aleixo, A., Aphalo, P. J., Asiegbu, F. O., Cabeza, M., Cairns, J., Candolin, U., Cardoso, P., Eronen, J. T., Hällfors, M., Hovatta, I., Juslén, A., Kovalchuk, A., Kulmuni, J., Kuula, L., Mäkipää, R., Ovaskainen, O., Pesonen, A.-K., … Vanhatalo, J. (2023). Understanding and applying biological resilience, from genes to ecosystems. Npj Biodiversity, 2(1), 1–13. 10.1038/s44185-023-00022-6

Tian, L., Preisser, E. L., Haynes, K. F., & Zhou, X. (2017). Social buffering in a eusocial invertebrate: Termite soldiers reduce the lethal impact of competitor cues on workers. *Ecology*, 98(4), 952–960. 10.1002/ecy.1746

Topaz, C. M., D’Orsogna, M. R., Edelstein-Keshet, L., & Bernoff, A. J. (2012). Locust Dynamics: Behavioral Phase Change and Swarming. PLOS Computational Biology, 8(8), e1002642. 10.1371/journal.pcbi.1002642

Townsend, S. W., Rasmussen, M., Clutton-Brock, T., & Manser, M. B. (2012). Flexible alarm calling in meerkats: The role of the social environment and predation urgency. Behavioral Ecology, 23(6), 1360–1364. 10.1093/beheco/ars129

Turetsky, K. M., Purdie-Greenaway, V., Cook, J. E., Curley, J. P., & Cohen, G. L. (2020). A psychological intervention strengthens students’ peer social networks and promotes persistence in STEM. Science Advances, 6(45), eaba9221. 10.1126/sciadv.aba9221

Vernier, C. L., Chin, I. M., Adu-Oppong, B., Krupp, J. J., Levine, J., Dantas, G., & Ben-Shahar, Y. (2020). The gut microbiome defines social group membership in honey bee colonies. Science Advances, 6(42), eabd3431. 10.1126/sciadv.abd3431

von Frisch, K. (1993). X. Variants of the “Language of the Bees.” In The Dance Language and Orientation of Bees (pp. 293–320). Harvard University Press. https://www.degruyterbrill.com/document/doi/10.4159/harvard.9780674418776.c41/html

Wasserman, S., & Faust, K. (1994). Social Network Analysis: Methods and Applications. Cambridge University Press.

Waters, J. S., & Fewell, J. H. (2012). Information Processing in Social Insect Networks. PLOS ONE, 7(7), e40337. 10.1371/journal.pone.0040337

Webster, M. M., & Ward, A. J. W. (2011). Personality and social context. Biological Reviews, *8c*(4), 759–773. 10.1111/j.1469-185X.2010.00169.x

Wey, T., Blumstein, D. T., Shen, W., & Jordán, F. (2008). Social network analysis of animal behaviour: A promising tool for the study of sociality. Animal Behaviour, 75(2), 333–344. 10.1016/j.anbehav.2007.06.020

Wild, B., Dormagen, D. M., Zachariae, A., Smith, M. L., Traynor, K. S., Brockmann, D., Couzin, I. D., & Landgraf, T. (2021). Social networks predict the life and death of honey bees. Nature Communications, 12(1), 1110. 10.1038/s41467-021-21212-5

Williams, T. D. (2007). Individual variation in endocrine systems: Moving beyond the ‘tyranny of the Golden Mean.’ Philosophical Transactions of the Royal Society B: Biological Sciences, 363(1497), 1687–1698. 10.1098/rstb.2007.0003

Wong, B. B. M., & Candolin, U. (2015). Behavioral responses to changing environments. Behavioral Ecology, 26(3), 665–673. 10.1093/beheco/aru183

Yamaguchi, Y., Ugajin, A., Utagawa, S., Nishimura, M., Hattori, M., & Ono, M. (2018). Double-edged heat: Honeybee participation in a hot defensive bee ball reduces life expectancy with an increased likelihood of engaging in future defense. Behavioral Ecology and Sociobiology, 72(8), 123. 10.1007/s00265-018-2545-z

Zhang, J., & Luo, Y. (2017). Degree Centrality, Betweenness Centrality, and Closeness Centrality in Social Network. Proceedings of the 2017 2nd International Conference on Modelling, Simulation and Applied Mathematics (MSAM2017). 2017 2nd International Conference on Modelling, Simulation and Applied Mathematics (MSAM2017). 10.2991/msam-17.2017.68

