## Supplemental Figures for "Behavioral compensation preserves collective behavior when individual members are compromised"

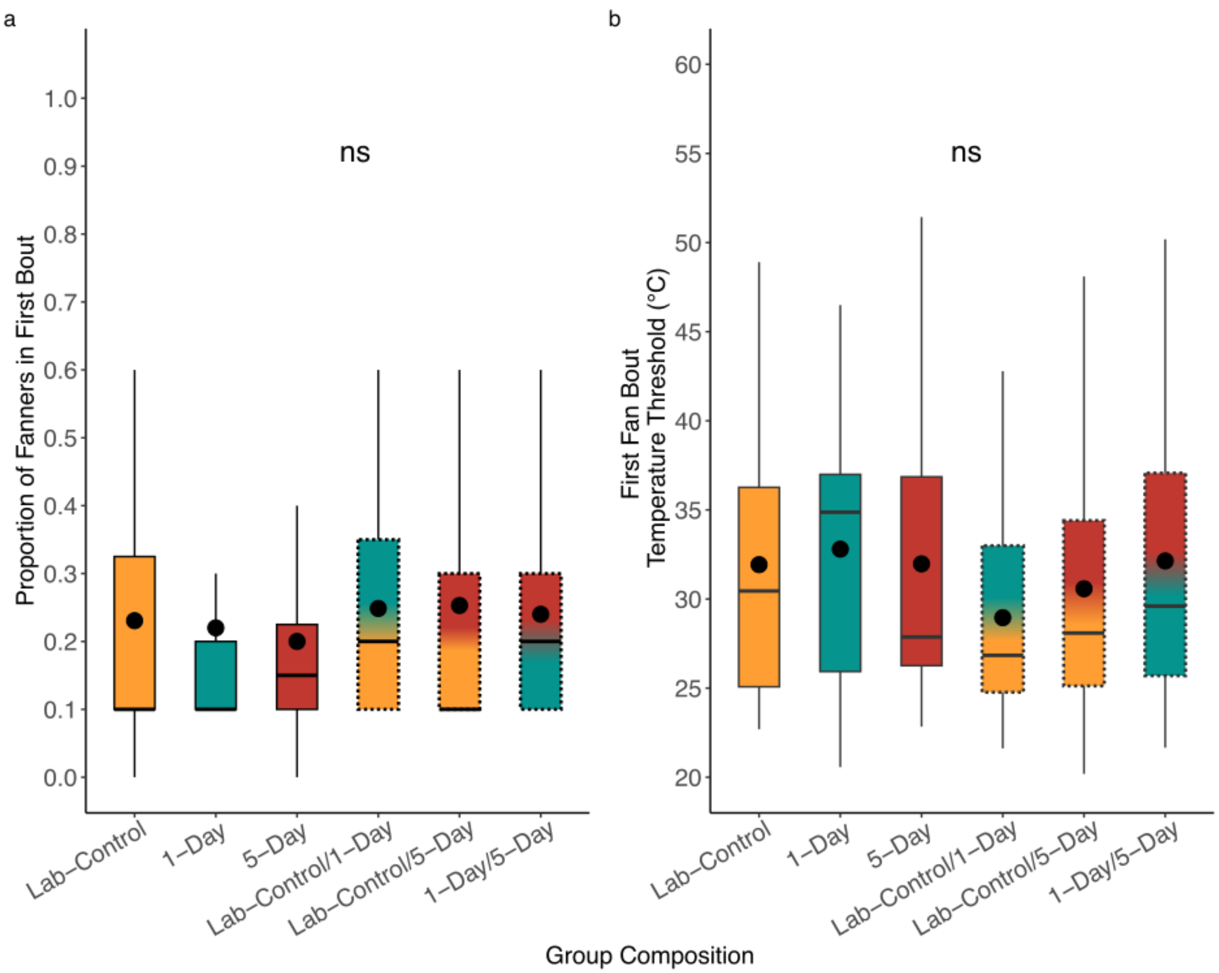

**Supplementary Figure 1. No statistical difference in proportion of fanners in the first bout and the temperature threshold the first bout of bees fanned at.** $N_{Lab\text{-}Control} = 38$, $n_{1\text{-}Day} = 39$, $n_{5\text{-}Day} = 35$, $n_{Lab\text{-}Control/5\text{-}Day} = 40$, $n_{Lab\text{-}Control/1\text{-}Day} = 36$, $n_{1\text{-}Day/5\text{-}Day} = 38$, for n = cages of 10 bees. First bout refers to the first group of fanners seen fanning in a trial. a) Proportion of fanners in the first bout of fanning seen in trials. No statistical difference was detected across all groups. GLM with a binomial distribution, $X^2 = 2.6567$, $p = 0.7527$ b) Temperature at which the first bout of bees was seen fanning at. No statistical difference was detected across all groups. LM, $X^2 = 2.5045$, $p = 0.7758$. Center black dots indicate mean while center lines indicate median. Letters indicate statistical significance.

**Supplementary Table 1.1 Max Fanning Bout Generalized Linear Mixed Model Estimates**

| | **Term** | **Estimate** | **Std. Error** | **z-value** | **Pr(>\|z\|)** |
|---|---|---|---|---|---|
| **Model Fixed Effects** | Intercept (LC) | 0.4627 | 0.2937 | 1.576 | 0.11512 |
| | 1D | -0.1908 | 0.1720 | -1.110 | 0.26714 |
| | 5D | -0.9345 | 0.1684 | -5.548 | <0.0001 |
| | LC/1D | -0.2841 | 0.1672 | -1.699 | 0.08928 |
| | LC/5D | -0.0239 | 0.1673 | -0.143 | 0.88644 |
| | 1D/5D | -0.5199 | 0.1690 | -3.077 | 0.00209 |
| | **Predictor** | **$X^2$** | **df** | **p-value** | |

| | | | | |
|---|---|---|---|---|
| **ANOVA Type II Wald Chi-Square Test** | Group Composition | 43.009 | 5 | **<0.0001*** ** |

**Supplementary Table 1.2 Max Fanning Bout Generalized Linear Mixed Model Tukey's Post-Hoc Test Results**

| Pairwise Comparison | Estimates | Standard Error | z-ratio | p-value |
|---|---|---|---|---|
| LC – 1D | 0.1908 | 0.172 | 1.110 | 0.8776 |
| LC – 5D | 0.9345 | 0.168 | 5.548 | **<0.0001*** ** |
| LC – LC/5D | 0.0239 | 0.167 | 0.143 | 1.000 |
| LC – LC/1D | 0.2841 | 0.167 | 1.699 | 0.5323 |
| LC – 1D/5D | 0.5199 | 0.169 | 3.077 | **0.0255** * |
| 1D – 5D | 0.7427 | 0.176 | 4.221 | **0.0003** *** |
| 1D – LC/5D | -0.1670 | 0.173 | -0.965 | 0.9291 |
| 1D – LC/1D | 0.0933 | 0.176 | 0.531 | 0.9949 |
| 1D – 1D/5D | 0.3291 | 0.176 | 1.875 | 0.4178 |
| 5D – LC/5D | -0.9106 | 0.172 | -5.297 | **<0.0001*** ** |
| 5D – LC/1D | -0.6504 | 0.170 | -3.827 | **0.0018** ** |
| 5D – 1D/5D | -0.4146 | 0.173 | -2.402 | 0.1553 |
| LC/5D – LC/1D | 0.2602 | 0.168 | 1.550 | 0.6315 |
| LC/5D – 1D/5D | 0.4960 | 0.167 | 2.965 | **0.0359** * |
| LC/1D – 1D/5D | 0.2358 | 0.167 | 1.415 | 0.7179 |

**Supplementary Table 1.3. Max Fanning Bout Generalized Linear Mixed Model Estimated Marginal Means Contrasts**

| Individual Treatment | Group Context | Probability of Fanning | SE | Estimated Marginal Mean | SE | df | z-ratio | p.value |
|---|---|---|---|---|---|---|---|---|
| LC | Uniform | 0.597 | 0.0657 | 0.268 | 0.170 | inf | 1.575 | 0.1152 |
| | Mixed | 0.660 | 0.0622 | | | | | |
| 1D | Uniform | 0.551 | 0.0679 | 0.175 | 0.179 | inf | 0.980 | 0.3270 |
| | Mixed | 0.594 | 0.0663 | | | | | |
| 5D | Uniform | **0.374** | 0.0644 | 0.521 | 0.170 | inf | 3.060 | **0.0022** * |
| | Mixed | **0.502** | 0.0681 | | | | | |

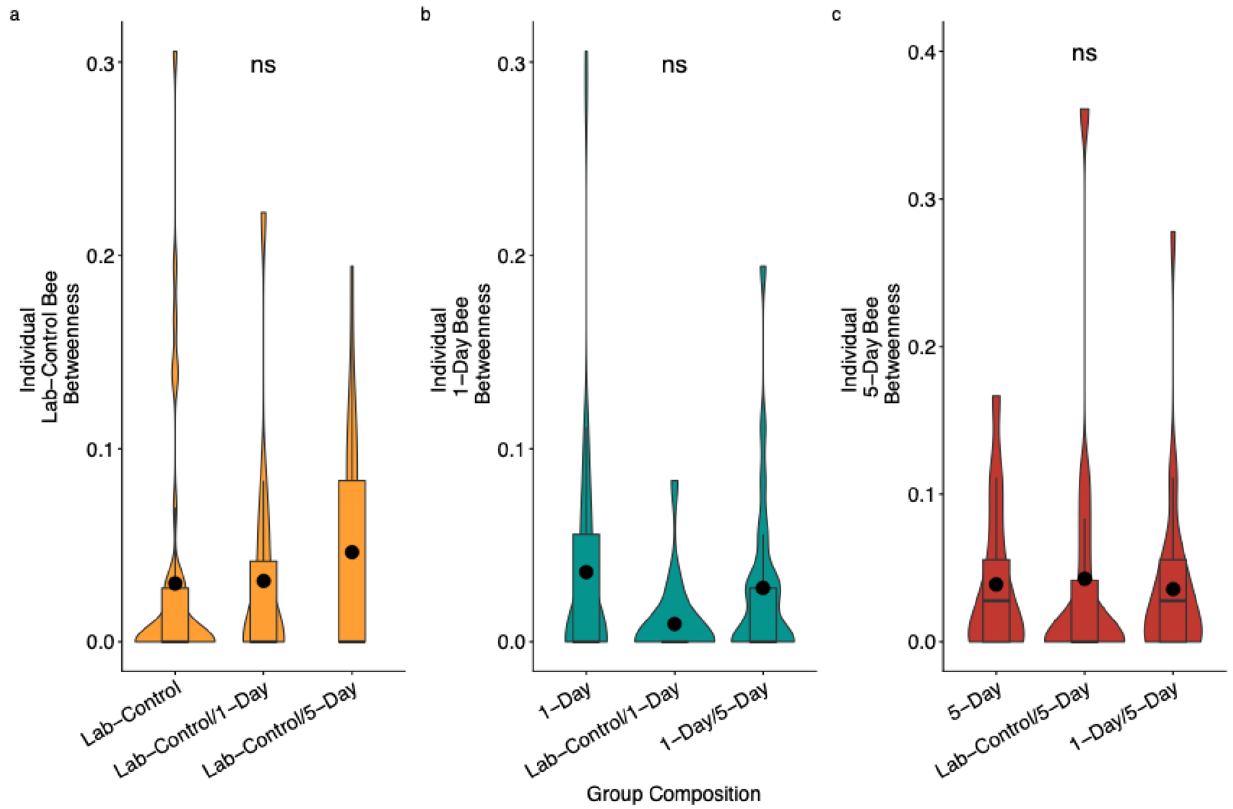

**Supplementary Figure 2. No statistical difference in individual betweenness across different group composition.** a) Lab control bees had no difference in betweenness regardless of group context. LM, F = 0.418, p = 0.6601 b) 1-Day bees had the same betweenness across group contexts. LM, F = 1.3047, p = 0.2772 c) 5-Day bees had no difference in betweenness metric across group context. LM, F = 0.0401, p= 0.9607. Center line indicates median, while center dots indicate mean. Letters indicate statistical significance where groups that do not share a letter are statistically different from one another.
